# Neuropixels Opto: Combining high-resolution electrophysiology and optogenetics

**DOI:** 10.1101/2025.02.04.636286

**Authors:** Anna Lakunina, Karolina Z Socha, Alexander Ladd, Anna J Bowen, Susu Chen, Jennifer Colonell, Anjal Doshi, Bill Karsh, Michael Krumin, Pavel Kulik, Anna Li, Pieter Neutens, John O’Callaghan, Meghan Olsen, Jan Putzeys, Charu Bai Reddy, Harrie AC Tilmans, Sara Vargas, Marleen Welkenhuysen, Zhiwen Ye, Michael Häusser, Christof Koch, Jonathan T. Ting, Neuropixels Opto Consortium, Barundeb Dutta, Timothy D Harris, Nicholas A Steinmetz, Karel Svoboda, Joshua H Siegle, Matteo Carandini

**Affiliations:** Allen Institute for Neural Dynamics, Seattle, WA, USA; UCL Institute of Ophthalmology, University College London, London, UK; Department of Neurobiology and Biophysics, University of Washington, Seattle, WA, USA; Janelia Research Campus, Howard Hughes Medical Institute, Ashburn, VA, USA; Department of Biomedical Engineering, Johns Hopkins University, Baltimore, MD, USA; IMEC, Leuven, Belgium; Wolfson Institute for Biomedical Research, University College London, London, UK; Allen Institute MindScope Program, Seattle, WA, USA; Allen Insti- tute for Brain Science, Seattle, WA, USA

## Abstract

High-resolution extracellular electrophysiology is the gold standard for recording spikes from distributed neural populations, and is especially powerful when combined with optogenetics for manipulation of specific cell types with high temporal resolution. We integrated these approaches into prototype Neuropixels Opto probes, which combine electronic and photonic circuits. These devices pack 960 electrical recording sites and two sets of 14 light emitters onto a 70 μm wide, 1 cm long shank, allowing spatially addressable optogenetic stimulation with blue and red light. In mouse cortex, Neuropixels Opto probes delivered high-quality recordings together with spatially addressable optogenetics, differentially activating or silencing neurons at distinct cortical depths. In mouse striatum and other deep structures, Neuropixels Opto probes delivered efficient optotagging, facilitating the identification of two cell types in parallel. Neuropixels Opto probes represent an unprecedented tool for recording, identifying, and manipulating neuronal populations.

## Introduction

Understanding brain function requires recording from myriad neurons, identifying them, and manipulating their activity. For large-scale recordings, an ideal method is extracellular electrophysiology via high-density electrodes such as Neuropixels probes^1,2^. For neuron identification^3-6^ and manipulation^7-11^, in turn, a leading method is optogenetics.

Electrophysiology and optogenetics are particularly powerful when paired with each other^12-14^. By combining them, one can test the causal role of specific neural populations by activating or inactivating those populations while recording the effects on neural activity^8,15-18^. One can also identify whether the recorded neurons belong to a genetic class of interest, by ‘optotagging^3-6^’: inducing this class to express an opsin and stimulating it with light. Optotagging is critical for connecting the wealth of knowledge about the gene expression, morphology, and connectivity of different cell classes to their function^19^.

Optogenetics, however, depends critically on delivering light with sufficient intensity and spatial resolution, potentially deep in the brain. This is difficult in brain tissue, which scatters and absorbs light^20^. It commonly requires inserting additional devices for light delivery, such as optical fibers^21-23^, waveguides^24^, or microLED arrays^25-29^. Existing approaches, however, have limited spatial resolution or light intensity, are invasive, and require a separate device for recording.

There is thus great interest in combining recording and light emission into a single ‘optrode’, but existing solutions have few recording sites or limited light intensity. Early optrodes integrated electrodes with optical fibers^30-34^, yielding few emitters. More emitters were enabled by micro-LEDs^28,35-39^. However, miniaturized microLEDs have low efficiency (1-3%) so even at moderate light they increase brain temperature^40,41^ by 0.5– 1.5 °C (Refs. ^25,28^). They thus deliver only low light intensities or duty ratios^42^.

To resolve these limitations, we combined Neuropixels recording technology with on-chip photonic waveguides that route high-intensity light down the shank of the probe. Light is generated outside the brain and routed by on-chip photonic waveguides^43-47^. This enables dual color illumination across a 1.4 mm span in parallel to voltage readout from close to a thousand selectable recording sites per shank with on-board amplification and digitization^1,2^. The resulting prototype device, called Neuropixels Opto, delivers unprecedented integration of high-resolution electrophysiology and optogenetics.

## Results

Neuropixels Opto integrates electronics and photonics to simultaneously record signals from 384 out of 960 recording sites and emit light from two sets of 14 emitters. The two sets of light emitters allow dual color optogenetics with blue and red light, making it possible to address two genetically-defined neural populations in parallel. For the red light, we chose a wavelength of 638 nm to excite highly effective red-sensitive opsins such as Chrimson^48^ and ChRmine^49^. This wavelength avoids the peaks of absorption by oxygenated blood^50^ at 420 nm and 540-580 nm and thus enhances the penetration of light into tissue. For the blue light, we set the wavelength to 450 nm (rather than the more common 473 nm) to efficiently activate Channelrhodopsin-2 (ChR2^51^) and its variants while reducing the activation of the red-sensitive opsins.

### Probe design

Neuropixels Opto probes monolithically integrate CMOS circuits for electrical recording with photonics for optical stimulation (**Fig. 1a,Suppl. Fig. S1**). An integrated silicon nitride (SiN) photonics layer provides photonic waveguides that route red and blue light to programmable emitters. The waveguides are fabricated in 150 nm thick SiN and are routed to the distal end of the shank. To couple out the light from the waveguides to the emitters and distribute it perpendicular to the probe, we used higher order, apodized Bragg grating couplers^52-55^ designed to spread the light over multiple diffraction peaks. The photonics layer lies above the CMOS platform designed for Neuropixels 1.0 probes^1^ (a 130-nm silicon-on-insulator CMOS Al process with 6 metal layers and Titanium nitride recording sites).

**Figure 1.**
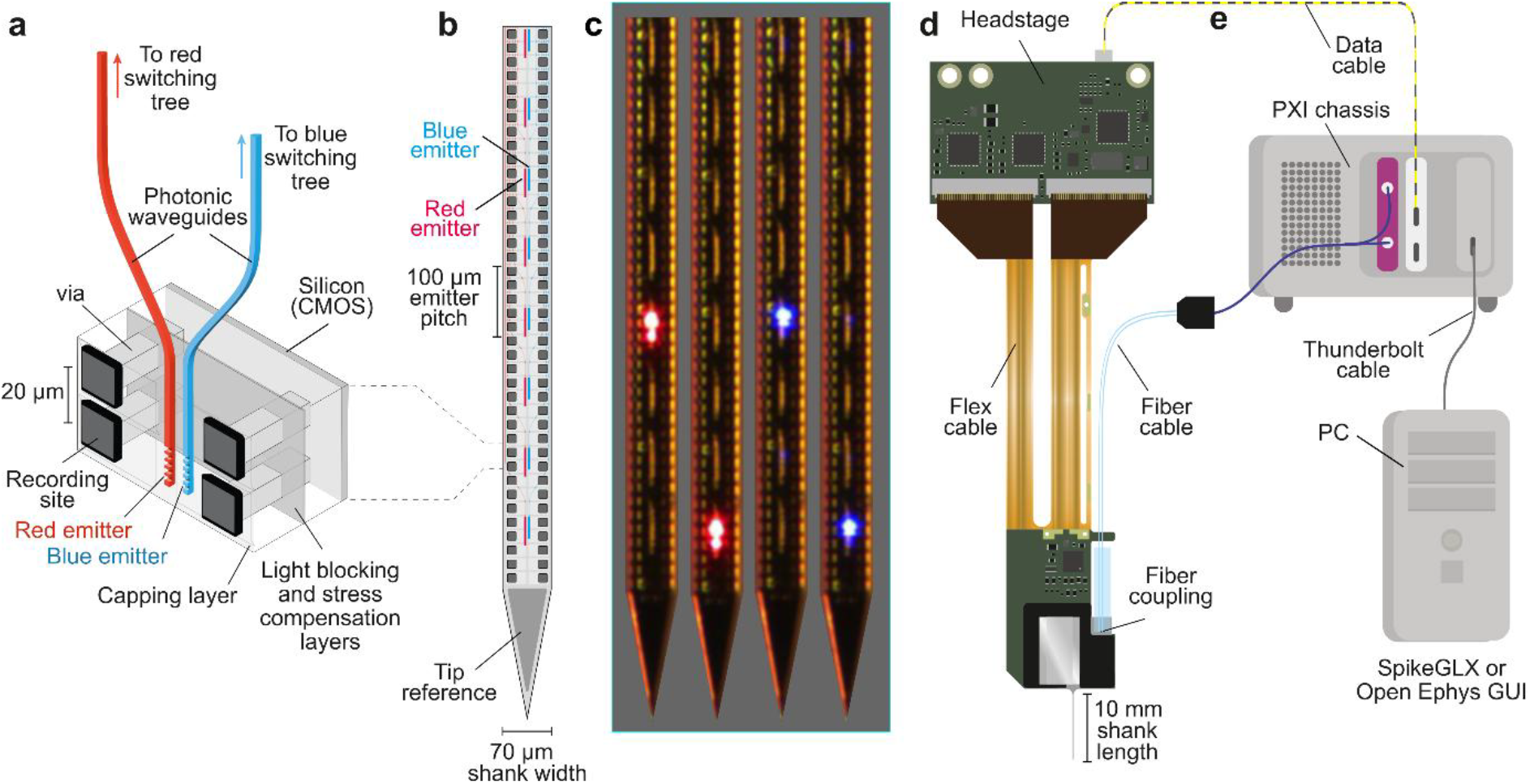
Design of the prototype Neuropixels Opto probe. **a**, Cross-section of the Neuropixels Opto shank, showing the titanium nitride (TiN) recording sites (connected with a “via” to the silicon CMOS layer) and the silicon nitride (SiN) photonic waveguides ending in the emitters (grating couplers). **b**, Layout of recording sites and dual color emitters. **c**, Photos of a probe shank across four time points, with two red and two blue emitters delivering light in succession. **d**, Device package. **e**, Neuropixels Opto system architecture, with PXI modules for data acquisition (*white*) and light delivery (*purple*).

This design posed two challenges. First, the addition of photonics can cause the probe shank to bend. We addressed this challenge by depositing a SiN compensation layer and a SiN capping layer (**Fig. 1a,Suppl. Fig. S1**), reducing tip deflection to < 200 μm. Second, scattered light from the photonic waveguides can interact with the CMOS circuitry, which is sensitive to light, increasing noise levels or introducing recording artifacts. To prevent light from reaching the CMOS circuits, we added a TiN/Al-based light blocking layer (**Fig. 1a,Suppl. Fig. S1**), keeping it as thin as possible to minimize shank bending and thickness.

The 14×2 emitters (16-25 μm^2^) are arranged on the center axis of the shank and are spaced 100 μm apart, covering 1.5 mm from the shank’s tip (**Fig. 1b,c**). The recording site array has a similar density to Neuropixels 1.0 (Ref. ^1^), with 960 titanium nitride (TiN) 12×12 μm^2^ recording sites separated vertically by 20 μm. The sites are arrayed in two vertical columns as in Neuropixels 2.0 (Ref ^2^), spaced 48 μm apart. The shank is 10 mm long, 70 μm wide, and 33 μm thick (9 μm thicker than 1.0 probes). As in 1.0 probes, signals from each recording site are split into bands for action potentials (AP, 0.3–10 kHz, digitized at 30 kHz) and local field potentials (LFP, <1 kHz, digitized at 2.5 kHz).

The light is generated by two fiber-coupled lasers at 450 and 638 nm, connected to the probe via grating couplers, and routed to the emitters by two photonic switching trees. There are 8 grating couplers: two to couple the light from the two fibers, and the others for active alignment of the fiber block and for measuring coupling losses. The light of each color is routed to the desired emitters via a programmable photonic binary switching tree (**Suppl. Fig. S2**). The switches are thermo-optic: Mach-Zehnder interferometers with thermal phase shifters based on the thermo-optic effect^56^. After calibration, they are the optical equivalent of a toggle switch. With four levels, the tree can specify 2^4^=16 outputs and thus address the 14 emitters independently. The current version of the probe allows one emitter per color to be on at a time, but any combination is possible in principle.

This switching tree was calibrated once, after fabrication. With high-intensity blue light, however, we encountered material instability, which resulted in fractions of light leaking from undesired emitters and would thus have required recalibration. We managed this issue by limiting the power of blue light. Therefore, in experiments requiring high light intensity and precise spatial addressing, we used red light.

The base of the probe integrates the fiber block, the photonics, and the CMOS recording circuits, which are mounted on a printed circuit board (PCB, **Fig. 1d**). To accommodate the fiber block and the photonics circuitry, we extended the 5-mm probe base integrating the recording circuits with two wings of 2 and 3 mm. The probe transmits data via flex cable to a headstage PCB, which connects to a digital data cable.

The data cable and the two-channel optical fiber cable are connected to two modules in a PXI base station, one for digital data processing and one containing blue and red lasers (**Fig. 1e**). Data acquisition, light delivery, and emitter selection are controlled by one of two widely used open-source software packages: SpikeGLX^1^ and Open Ephys GUI^57^, which were updated for the purpose.

### Electrical and optical characterization

Despite the addition of the photonics, the electrical performance of the Neuropixels Opto probe remained similar to the widely used Neuropixels 1.0 probes^1^. The average electrode impedance was 138 ± 27 kΩ, and the average RMS noise in the AP and LFP bands was 5.45 ± 0.02 μV and 5.33 ± 0.03 μV (mean ± s.e., N = 20,097 based on 957 sites from 21 probes, **Suppl. Fig. S3**). These values are below the typical measurements in Neuropixels 1.0 (5.5 μV in AP band and 8.0 μV in LFP band^1^) and Neuropixels 2.0 (7.2 μV for the combined AP-LFP band^2^). This low noise, combined with high site density (100 sites/mm), enables recordings with similar quality as those routinely obtained with established Neuropixels probes.

As expected, the light at the emitters was a small fraction of the light delivered by the optical fibers: the average emitted power was 2.07 ± 0.02 % of the input power for red light, and 0.24 ± 0.01 % for blue light (mean ± s.e., N = 434 based on 14 emitters from 31 probes, **Fig. 2a**). The attenuation (– 16.99 ± 0.06 dB for red, –26.47 ± 0.08 dB for blue) is caused by the coupling of fiber to waveguide (∼6 dB), the switching tree (∼4 dB), the emitter (simulated < 4 dB) and waveguide propagation, where losses are particularly strong with blue Neuropixels Opto light (3.0 dB/cm, compared to 0.5 dB/cm for red light). Because of these losses, achieving 100 μW of output at the emitters requires input powers of ∼5 mW for the red light and ∼40 mW for the blue light. These powers are easily delivered by external lasers.

**Figure 2.**
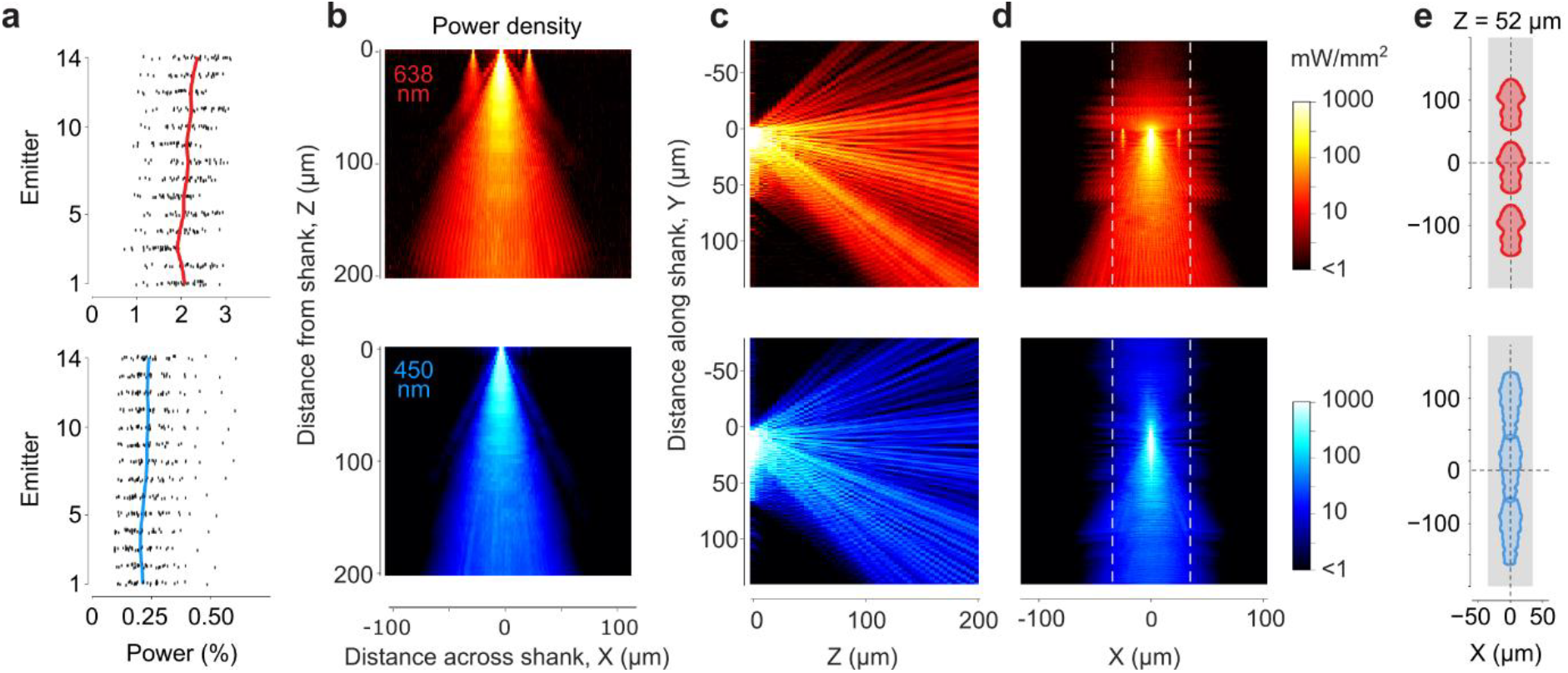
Optical characterization of the Neuropixels Opto probe. **a**, Efficiency of the emitters, showing output light power as a percentage of input power for 14 red (*top*) and blue (*bottom*) emitters from N = 31 probes. Each dot corresponds to one emitter from one probe. Curves show average over probes. **b**, Top view, showing light propagation from a red and blue emitter, measured in water. The color (color scale in panel d) indicates the max projection. These measurements were made on a test structure where emitters were placed 25 μm apart rather than the 100 μm of the prototype probe (Methods) and this led to small imaging artifacts visible in the red emission (*top*), where two emitters to the side of the central one also appear to emit light. **c**, Same data, projected over a side view. **d**, Same data, projected over a front view. Dashed lines delineate the width of the probe. **e**, Section on a plane located 52 μm away from the shank, showing areas where power density is >10 mW/mm^2^ (for a 100 μW output) for three nearby emitters.

The emitted light is sufficient for optogenetic manipulations of neurons in the vicinity of the probe. For ChR2, estimates of the light intensity required for optogenetic stimulation range from 10 mW/mm^2^ (Ref. ^51^) to 5 mW/mm^2^ (Ref. ^58^) to 1 mW/mm^2^ (Ref. ^59^). The variability perhaps reflects differences in preparations, opsin expression levels and distributions, and spatial overlap of excitation light with the neuron’s membrane. To be conservative we used the higher estimates, and measured the volume where light power density is at least 10 mW/mm^2^. For a 100 μW output, this volume exceeds 470,000 μm^3^, extending >100 μm from the shank over a wide angular range (**Fig. 2b-d** and Suppl. Movie). This is more than sufficient to stimulate neurons in the vicinity of the recording sites and beyond the ∼50 μm limit for single-unit recordings^60,61^. The volume where light power density is >1 mW/mm^2^ is considerably larger, and exceeds the range of the microscope used for these measurements. Moreover, in a scattering medium such as the brain, this volume is expected to be more homogenous than in water. We confirmed this expectation using an optical phantom matched to the scattering observed in rodent gray matter^62^ (**Suppl. Fig. S4**).

The close spacing of the emitters means that there is a minimal gap between the patterns of light emitted by different emitters: when slicing the emission profile close to the shank (in a plane ∼50 μm away), the area with a power density >10 mW/mm^2^ largely tiles the shank axis (**Fig. 2e**).

By delivering light away from the recording sites, Neuropixels Opto probes largely avoid photoelectric artifacts. Direct illumination of Neuropixels recording sites with sharp-onset surface stimulation creates a photoelectric artifact in the range of 1 mV or more^1^. By contrast, illumination via Neuropixels Opto emitters, which point light away from the recording sites, caused only a small electrical artifact of ∼30 μV. This artifact was present only with red light, and only with sharp onsets. It was uniform across recording sites and thus easily corrected with standard preprocessing steps (**Suppl. Fig. S5**).

### Activating local neural populations

We next established the ability of the probes to activate spatially separated neuronal populations. We inserted Neuropixels Opto probes acutely in the primary visual cortex of awake, head-fixed mice following local viral expression of the red-sensitive depolarizing opsin ChRmine^49^ under the CaMK2 promoter (**Fig. 3a,Suppl. Fig. S6**), which preferentially^63^ targets excitatory neurons. Cells expressing CaMK2 are largely (though not uniquely^64-67^) excitatory. Tapered pulses of red light (638 nm) lasting 400 ms at one example emitter elicited neural activity that was restricted to recording sites near the emitter (**Fig. 3b**). These recordings had similar quality to standard Neuropixels probes^1,2^: they yielded 0.23±0.09 units/site (median ± m.a.d., N = 13 recordings in 3 mice), similar to the yield obtained in the same area in a set of recordings^68^ with Neuropixels 1.0 probes (0.22±0.10 units/site, median ± m.a.d., N = 20 recordings in 20 mice in 10 labs, filtered with the same quality metrics as our study). We could then readily spike-sort them to obtain the spikes of individual neurons (**Fig. 3c**), which were unaffected by optical stimulation (**Suppl. Fig. S7**).

**Figure 3.**
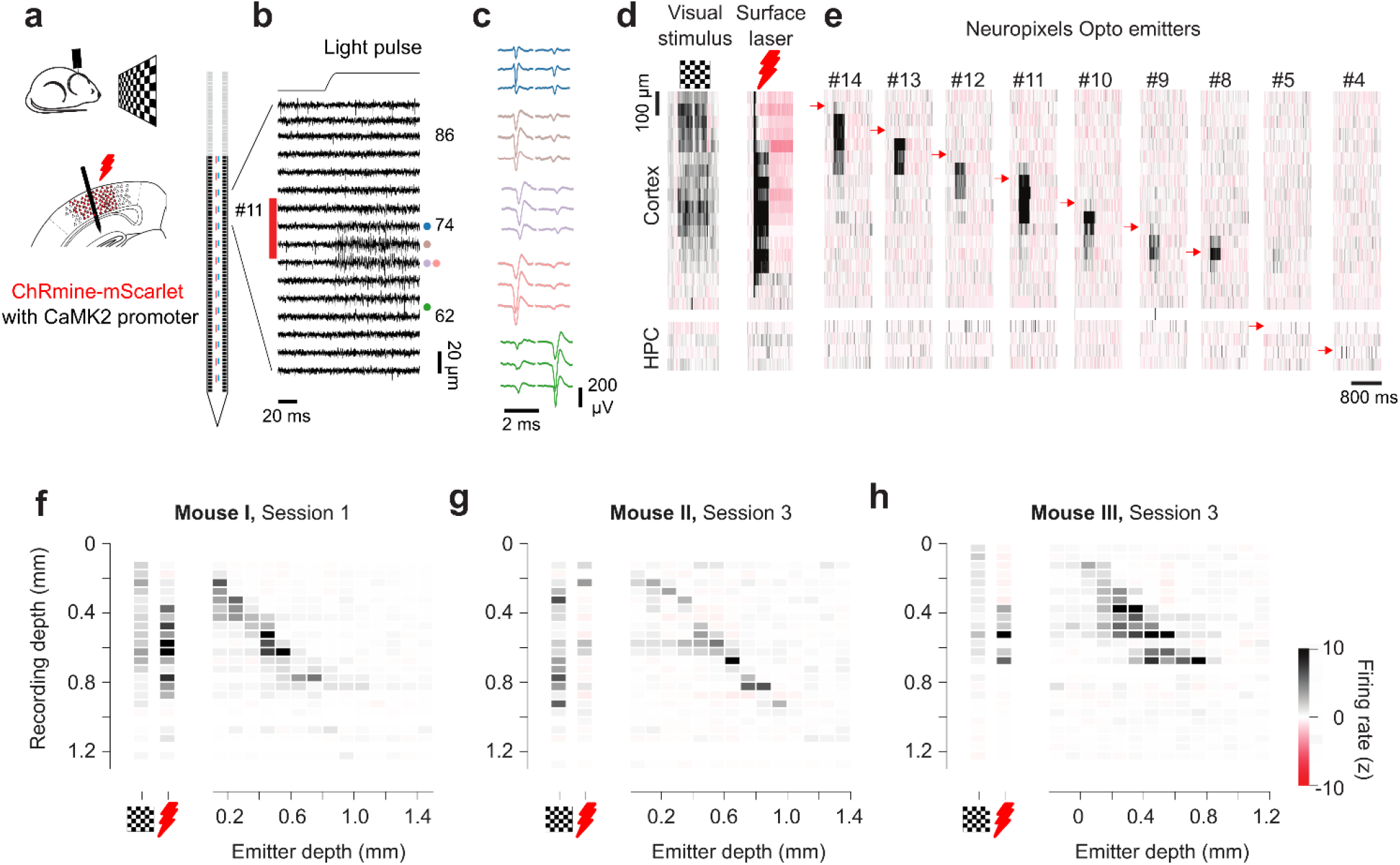
Using Neuropixels Opto to record and activate localized neural populations. **a**. We inserted a Neuropixels Opto probe ∼1.4 mm deep in the visual cortex of mice expressing the red-sensitive opsin ChRmine (conjugated with mScarlet) in cortical neurons under the CaMK2 promoter. Mice viewed a visual stimulus, and an additional red laser could illuminate the surface of the posterior cortex. **b**. Simultaneous Neuropixels Opto recordings and optical stimulation with an example emitter (#11). Recordings (while the screen was gray, with no external laser), show baseline activity 50 ms prior to emitter photostimulation and strong spiking activity after stimulation onset on a subset of recordings sites near emitter 11. **c**. Average spike waveforms from five example single units recorded on sites near emitter 11. The mean waveform was calculated across 100 spikes. **d**. Average firing rate (bin size 50 µm x 2 ms, 40 trials) for the same recording session, plotted as a function of depth, showing cortical responses to visual stimulation and surface illumination. Color scale bar appears at the bottom right of the figure. **e**. Responses of the same neurons to single emitter activations at different depths (*arrows*). **f**. Summary of these data showing average over time of response during stimulation with visual stimulus, surface laser, and single emitters (abscissa), at different cortical depths (ordinate). **g-h**. Same format, for example insertions in two other mice. In different insertions the emitters were at different cortical depths. In **h**, the top emitter was outside the cortex (negative depth), where it elicited no activity. Additional measurements in these mice are shown in **Suppl. Fig. S8**.

To establish baseline measurements, we presented a visual stimulus (full-field checkerboard) and we illuminated the surface of the posterior cortex with a red laser (638 nm, 5 mW). These baseline measurements indicated the presence of recordable neurons throughout the depth of the cortex. We summarize their activity in terms of firing rate as a function of cortical depth (**Fig. 3d**).

By activating one emitter at a time, Neuropixels Opto probes could spatially address different subpopulations of these neurons with high resolution (**Fig. 3e**). Trials with stimulation from different emitters were randomized and randomly interleaved with the baseline trials (visual stimulus or surface laser). Stimulation by single emitters activated smaller groups of these neurons at nearby depths. Taking the average over time of these responses revealed an approximately diagonal matrix (**Fig. 3f**), reflecting concordance between the location of stimulation and the location of neurons with increased firing rates.

Similar results were obtained in two other mice (**Fig. 3g,h**) and across multiple experiments (13 probe insertions in 3 mice, **Suppl. Fig. S8**). At the light intensities provided in these experiments, the population of activated neurons extended vertically over 151±71 μm (full width at half maximum, mean ± s.d., N = 13 insertions). This extent represents a lower bound, because the recorded neurons are likely to be a fraction of the activated ones. It is comparable to the 100 µm spacing between emitters, suggesting that the probe can provide spatial coverage across emitters. Measurements of local field potential and current source density confirmed that the emitters generated localized current sources and sinks (**Suppl. Fig. S9**).

To exclude possible artifacts of light stimulation due to thermal effects^69^ or to activation of the retina^70^ we performed two sets of measurements. First, we confirmed that the emitters did not evoke activity in a region that did not express the opsin: the hippocampus (**Fig. 3e-h**). Second, we ran a control experiment in a mouse that received no virus injection, and found no effect of light stimulation (**Suppl. Fig. S10**).

These results indicate that Neuropixels Opto probes provide concurrent large-scale recordings and fine spatially addressed optogenetics across the depth of a brain structure.

### Driving localized circuits

We next tested the ability of Neuropixels Opto probes to drive local circuit effects such as those mediated by synaptic inhibition. We expressed the red-sensitive depolarizing opsin ChrimsonR^48^ in putative inhibitory forebrain neurons by systemic injection^71^ of a DLX2.0 enhancer virus^72-74^ (**Suppl. Fig. S11**). We then inserted Neuropixels Opto probes in the dorsal cortex of awake, head-fixed mice, and we delivered 250 ms pulses of light at random times from random emitters.

The results were consistent with localized optogenetic activation of inhibitory neurons, and consequent synaptic inhibition of excitatory neurons. Light delivery activated some neurons and inactivated others (**Fig. 4a-d**). Because the opsin depolarizes neurons in which it is expressed, the activated neurons should correspond to the putative inhibitory neurons expressing the opsin, while inactivated neurons could only reflect neurons receiving synaptic inhibition from the activated population. To test this interpretation, we analyzed the spike waveforms^75^ and distinguished putative fast-spiking neurons, which have narrow spikes, from the rest of the neurons, which are likely to be mostly pyramidal and have broader spikes^76^. As expected^75^, the activated neurons were predominantly fast-spiking, whereas the putative pyramidal neurons were predominantly inactivated (**Fig. 4e**). Moreover, cross-correlograms^76,77^ between some pairs of activated and inactivated neurons were consistent with putative monosynaptic inhibitory connections (**Fig. 4f**). Both activated and inactivated neurons were observed primarily at depths near the emitter (**Fig. 4g**), indicating that the localized activation of the activated putative inhibitory neurons engaged localized circuit effects. These results were replicated in 10 sessions in 3 mice, providing consistent results (**Fig. 4e**,**g** and **Suppl. Fig. S12**). Both the activation of putative inhibitory neurons and the resulting inactivation of putative excitatory neurons grew in strength with increasing light intensity, especially at low intensities (**Suppl. Fig. S13)**. Taken together, these results confirm that Neuropixels Opto probes are suitable for causing spatially-localized circuit effects mediated by synaptic transmission.

**Figure 4.**
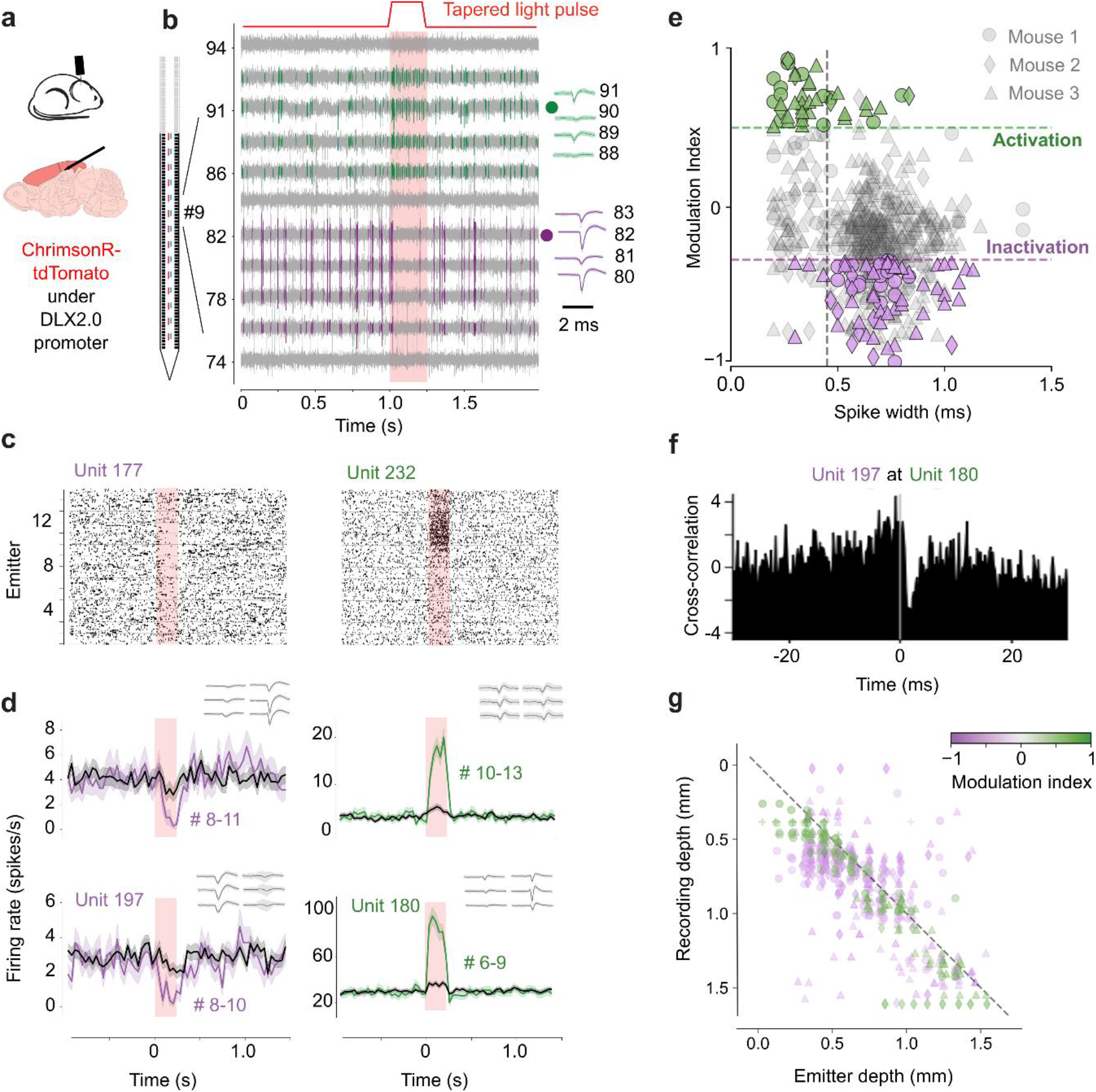
Using Neuropixels Opto to drive localized circuits. **a**. We inserted a Neuropixels Opto probe in the dorsal cortex of mice expressing red-sensitive depolarizing opsin ChrimsonR-tdTomato in putative inhibitory neurons (via DLX2.0 enhancer virus^72-74^). **b**. Electrical signals at 20 recording sites during red light stimulation from emitter 9. Colors indicate the spikes of two nearby units, one activated (unit 189, *green*) and one inactivated (unit 208, *purple*) by light. Waveforms across peak channels (*right*) confirm neural activity. In this panel and subsequent ones, the shaded rectangle indicates the time of optical activation. **c**. Spike rasters for a pair of example units, one inactivated (unit 177) and one activated (unit 232) by light, ordered by the stimulating emitter (*ordinate*). **d**. Average firing rate (40 ms bin size) relative to light onset for those units and for two additional units (197 and 180). Shading indicates ±1 s.e. Waveforms (*insets*) show spike shapes across 6 peak recordings sites. **e**. Spike width (trough to peak) of average waveforms vs. effect of light stimulation, measured by a modulation index (R_1_-R_0_)/(R_1_+R_0_) where R_0_ and R_1_ are firing rates before and during stimulus, showing significantly activated units (*green*) and inactivated units (*purple*) at p < 0.005 (paired t-test). Narrow spikes were defined as width < 0.4 ms (*vertical line*). **f**. Crosscorrelogram between units 197 and 180. **g**. Recording vs. emitter depth for significantly modulated neurons. Each neuron appears at one or more emitter depths (if modulated by light from multiple emitters).

### Optotagging nearby neurons

When light-sensitive opsins are expressed in a cell-type-specific manner, these cells can be identified in extracellular recordings via optotagging^3-5^, i.e. based on their low-latency responses to pulses of light. An ideal tool for optotagging would have minimal artifacts in response to light pulses, minimal activation of neurons outside the range of recording, and the ability to deliver both blue and red light wavelengths deep inside the brain. Neuropixels Opto probes meet all of these criteria. As we have seen, moreover, by directing the light away from the recording sites, the probes largely avoid photoelectric artifacts (**Suppl. Fig. S5**). Indeed, the spike waveforms occurring during light stimulation had similar shape to those occurring when no stimulation was present (**Suppl. Fig. S14**).

To test the efficacy of Neuropixels Opto for optotagging two cell types in parallel, we recorded in mice expressing a blue-sensitive opsin in one population and a red-sensitive opsin in a second population. Neurons expressing blue-sensitive opsins, such as CoChR^48^, will only be depolarized by blue light, as these opsins’ sensitivity curves drop sharply at longer wavelengths, making them largely unresponsive to red light. By contrast, neurons expressing red-sensitive opsins, such as ChRmine^49^, will be depolarized by both blue and red light, due to these opsins’ long tail of sensitivity to shorter wavelengths. These distinct response patterns make it possible to identify two cell types in the same experiment.

The red-shifted opsins were expressed with Cre driver lines and Cre-dependent AAVs, whereas the blue-shifted opsin (CoChR) was expressed with enhancer AAVs^78^. We injected an enhancer AAV in the striatum to express CoChR-EGFP in direct-pathway (D1) medium spiny neurons (MSNs), indirect-pathway (D2) medium spiny neurons, or cholinergic (Chol) interneurons (**Suppl. Fig. S15**). *Ex vivo* slice recordings revealed peak photocurrents of up to 6 nA in CoChR-EGFP+ cells, with mean peak photocurrents of 3.2 nA for D1 neurons, 3.0 nA for D2 neurons, and 3.3 nA for Chol neurons (**Suppl. Fig. S16**). These currents were significantly higher than obtained in D1 neurons expressing ChR2(H134R) (Ref. ^79^), which had a mean peak photocurrent of 0.4 nA (P < 0.001, Welch’s one-way ANOVA with post-hoc t-tests corrected for multiple comparisons). For *in vivo* experiments, we thus chose CoChR for its higher blue light sensitivity and photocurrents. The redshifted opsins ChrimsonR and ChRmine were selected for similar reasons.

To optotag neurons we delivered sequences of 10 ms light pulses^5,6^ at 20 Hz. Previous work^6^ has shown that low-latency responses to light pulses in quick succession are only seen in neurons that are activated directly by light (because they are not abolished by synaptic blocker^6^). We thus considered neurons to be optotagged if they satisfied three criteria: (1) they displayed a significant increase in firing rate compared to baseline in response to a minimum of 4 out of five 10 ms pulses from at least one emitter; (2) the driving emitter evoked at least one spike in 30% of the trials; (3) the latency of the evoked spikes was < 8 ms.

To illustrate this approach, consider an example experiment where we expressed the blue-sensitive opsin CoChR-EGFP^48^ in D1 MSNs with an enhancer AAV^78^, and the red-sensitive opsin ChRmine-mScarlet^49^ in D2 MSNs via a Cre-dependent AAV in a specific (Adora2a-Cre) driver line (**Fig. 5a**, see Methods for virus details). Because the photoartifacts were small and disappeared after preprocessing (**Suppl. Fig. S5**), spikes were readily identifiable in the raw traces around each light pulse (**Fig. 5b**). For example, consider two units tagged by red light pulses from emitter 3 (**Fig. 5c**). Each unit shows consistent, low-latency spiking responses to each of five 100 µW pulses across 50 trials (**Fig. 5d**). The strongest responses were evoked by the emitter near the estimated position of the soma (**Fig. 5e**). For units tagged with blue light, some longer-latency spikes were evoked also by emitters distant from the soma, perhaps due to the small amounts of light leakage mentioned earlier (which are specific to blue light and would require recalibration to be removed). In this recording, we were able to tag most of the recorded units in striatum (25 out of 39, **Fig. 5f**), allowing direct comparison of the activity of populations of two cell types in a single structure.

**Figure 5.**
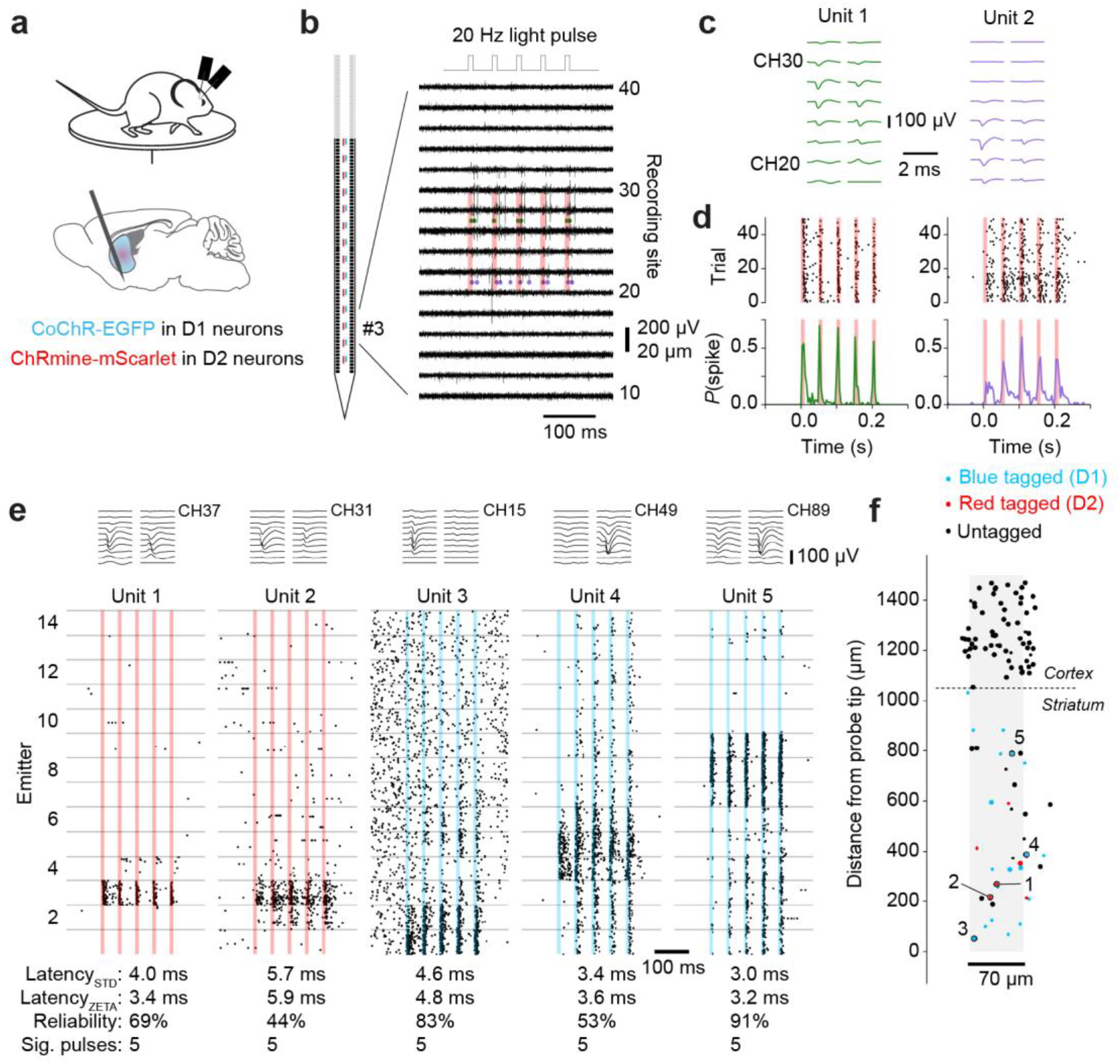
Using Neuropixels Opto for optotagging. **a**. We inserted two Neuropixels Opto probes in the striatum of Adora2a-Cre mice expressing the blue-sensitive opsin CoChR in D1 MSNs (via a D1-MSN-specific enhancer virus) and the red-sensitive opsin ChRmine in D2 MSNs (via a Cre-dependent AAV). Mice were free to run on a disc. After 20 min of recording we ran the optotagging protocol (10 ms, 20 Hz, 100 µW pulses from each of 14 blue or red emitters, randomly interleaved). **b**. Recorded traces for one trial of light presentation, showing spike times (green and red dots) of two example units tagged by red light from emitter 3. Red shaded areas indicate the timing of the light pulses. **c**. Mean waveforms for the two units highlighted in panel b. **d**. Spike raster and peri-stimulus time histogram for 50 trials of stimulation from emitter 3, showing consistent, low-latency response to each light pulse. **e**. Stacked rasters across 50 trials from all 14 emitters for five example units (including the two units from panels b–d). Red and blue shaded regions indicate the time of light presentation. Mean waveforms are shown above each raster. Key optotagging metrics are listed below each raster; units are considered tagged if they have a median latency below 8 ms, a reliability above 30%, and a significant response to at least 4 pulses. Latencies calculated via the ZETA test^82^ are shown for comparison. **f**. Estimated location of all units passing quality control from a single recording, with units activated by blue or red light shown in blue and units activated by red light only shown in red. Large dots indicate units that pass quality metric thresholds for the complete session (ISI violations ratio < 0.5, amplitude cutoff < 0.1, presence ratio > 0.8). Small dots indicate units that pass the ISI violations ratio threshold only for the pre-stimulus baseline interval (typical of units with low spontaneous firing rates that are strongly driven by light presentation).

Similar results were obtained in multiple other experiments where we used a variety of expression strategies for parallel optotagging of D1 and D2 MSNs (as in the example above), or D2 MSNs and Chol interneurons, or D1 MSNs and Chol interneurons, or for tagging glutamatergic or GABAergic neurons in the midbrain (**Suppl. Fig. S18**).

Overall, we optotagged 261 units with Neuropixels Opto across 40 sessions in 26 mice, using CoChR^48^ and ChRmine^49^, ChrimsonR^48^, rsChRmine^80^, or somBiPOLES^81^ (**Suppl. Table 1**).

We combined the results of these experiments to assess whether optotagging was equally possible at all distances from an emitter or whether there were gaps in coverage. We estimated the 2D location of every unit based on the spatial distribution of its spike waveform, and compared the locations of optotagged and untagged units (**Fig. 6a,b**). We first expressed each unit’s position relative to the driving emitter (the emitter evoking the largest response), and found that optotagged units tended to be located below the driving emitter (i.e., closer to the probe tip, **Fig. 6c**, p < 1e–10, Wilcoxon signed-rank test). This arrangement is consistent with the illumination profile of the emitters, which direct light downward along the shank’s long axis (**Fig. 2e)**. We then calculated each unit’s position relative to the nearest emitter, so that we could inspect the distribution of tagged neurons in the horizontal and vertical dimensions (**Fig. 6d**). Horizontally, the tagged units spanned the entire 70 µm width of the shank, with a slightly higher density (than untagged units) near the center of the shank (**Fig. 6e**, p = 0.0036, Kolmogorov-Smirnov test). Vertically, i.e. along the length of the shank, instead, the distribution of tagged units was indistinguishable from that of untagged units (**Fig. 6f**, p = 0.58, Kolmogorov-Smirnov test). Thus, the 100 µm inter-emitter spacing is sufficiently dense to leave no gaps in optotagging coverage.

**Figure 6.**
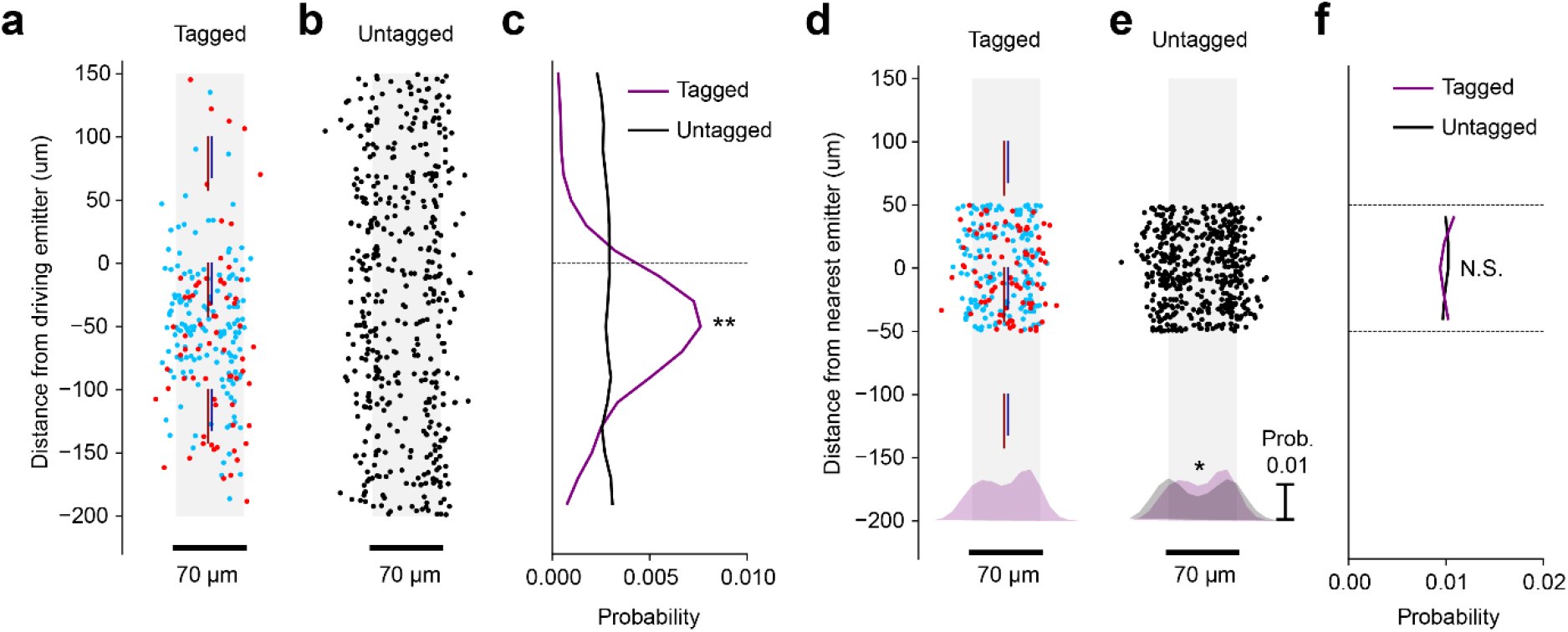
Spatial distribution of optotagged units. **a**. Estimated location of units relative to the driving emitter, for units that were optotagged by red light (*red dots, N* = 83 units from 25 sessions) and blue light (*blue dots, N* = 178 units from 26 sessions). **b**. Same as panel a, but for units from the same sessions that were not optotagged (*black dots, N* = 500 randomly selected units from 40 sessions). For these untagged units, the driving emitter is selected arbitrarily. **c**. Probability of tagging a unit as a function of distance from the driving emitter along the shank insertion axis. Most tagged units are located below the driving emitter (deeper in the brain). ** = *P* < 1e–10, Wilcoxon signed-rank test for difference from zero. **d-f**. Same data as panels a-c, plotted relative to the nearest emitter (rather than the driving emitter), showing that units have a similar probability of being optotagged at any location within 50 μm of an emitter. The histograms at the bottom of d and e show the distributions of unit locations orthogonal to the shank insertion axis. * = *P* < 0.01, N.S. = *P* > 0.05, Kolmogorov–Smirnov test for equality of distributions. See **Suppl. Table 1** for information about cell types and opsins in these experiments.

## Discussion

Thanks to integrated CMOS and photonics, Neuropixels Opto probes provide a single device for large-scale neural recordings and spatially addressable optogenetics. Our tests demonstrate that these probes can precisely manipulate neural activity near emitters in the intact brain. They thus deliver fine spatially-addressable optogenetics across the depth of a brain structure, while providing high-resolution, large-scale recordings. This capability is unprecedented, and is ideal for investigating the circuit organization of the cerebral cortex^8,15-18^ and other brain regions. In addition, Neuropixels Opto is well-suited for optotagging^3-5^, making it possible, for the first time, to identify the cell type of the majority of units in regions such as the striatum.

Our measurements in the cerebral cortex demonstrated that Neuropixels Opto probes can elicit highly localized optogenetic activity. Indeed, the effects that we observed (**Fig. 3**) are remarkably localized given that the layers of the cortex are highly interconnected^15^ through both direct cortical connections and indirect thalamic pathways. Indeed, previous studies that activated individual cortical layers optogenetically have found enhancement^83-85^ or suppression^86-88^ of activity in other layers. The highly localized results obtained with Neuropixels Opto might be due to lower light levels and to the focused range of the probe’s emitters.

Nonetheless, the spatial resolution obtainable with Neuropixels Opto is limited by the fundamental constraints of one-photon optics in a scattering medium, and by the morphology of neurons and opsin expression within those neurons. The scattering of light in brain tissue (simulated in **Suppl. Fig. S4**) is such that no emitter can fully restrict activation to highly localized volumes. Moreover, the region of activation can be further increased by the morphology of neurons and the distribution of opsin expression in extended neural processes. For example, the activation of neurons tens of µm below the emitter (**Fig. 6**) is likely a combined result of light scattering and distributed opsin expression. In this respect, future experiments may obtain more localized activations by using opsins that target specific portions of the neurons, such as the soma or the axon initial segment^89^.

Neuropixels Opto probes represent a marked technical achievement, as they are substantially more complex to fabricate than the existing Neuropixels 1.0 and 2.0 probes. This complexity can be measured as the number of processing steps required for layer depositions, patterning of the layers, and quality inspection. The prototype Neuropixels Opto probes presented here require ∼740 such processing steps, almost twice as many as for the Neuropixels 1.0 and 2.0 probes, which require ∼400 processing steps.

As with Neuropixels 1.0 and 2.0 probes^1,2^, our model is to produce the Neuropixels Opto probes in quantity and distribute them at cost to a wide community. Here we have demonstrated successful prototype probes. Turning these prototype probes into a mass-producible probe requires additional rounds of fabrication and testing. As with the 1.0 and 2.0 probes, during this process we will seek to further adjust the design in multiple ways. First, we aim to make the bluelight switches and waveguides more robust by putting them in a separate photonic layer. Second, we aim to integrate photodetectors to monitor the power of each emitter, thus providing active feedback and the ability for users to recalibrate the optical switches if any cross-talk is observed across emitters. Third, we aim to simplify the coupling between lasers and probe, and reduce the probe’s form factor, by upgrading the CMOS backend to the more compact design developed for the 2.0 probes^2^. Fourth, we aim to further increase the number of red and blue emitters.

We anticipate that Neuropixels Opto probes will become an essential tool for combining high-density electrophysiological recordings with local optogenetic activation or inactivation, and for cell type-specific electrophysiology across the brain.

## Acknowledgments

We thank numerous colleagues who contributed to this project. At IMEC, Enrico Tonon contributed invaluable assistance. At UCL, Bex Terry and Magdalena Robacha assisted with colony management and brain sectioning. At UW, Ljuvica Kolich and Kimberly Miller assisted with mouse husbandry. At AI, the Allen Institute Transgenic Colony Management, Lab Animal Services, Neurosurgery & Behavior Team, and Viral Core provided invaluable assistance, Ximena Opitz-Araya managed cloning of viral vectors, and Alessio Buccino helped with spike sorting pipelines. At HHMI, John Macklin provided advice on optical phantoms. This research was funded by the Wellcome Trust (grants 204915/Z/16/Z to MC, MH, and TDH and 225819/Z/22/Z to MC, MH, and NAS), the Howard Hughes Medical Institute (TDH), the BRAIN Initiative (U01NS133760 to JHS and KS; UF1MH128339 to JT), the Paul G Allen Family Foundation (CK, JHS and KS), the Research Council of Norway (NORBRAIN 350201 to EIM), the Pew Biomedical Scholars Program (NAS), and the Klingenstein-Simons Fellowship in Neuroscience (NAS). MC holds the GlaxoSmithKline / Fight for Sight Chair in Visual Neuroscience.

## Consortium membership

The Neuropixels Opto Consortium includes Matteo Carandini (UCL), Barun Dutta (IMEC), Timothy D Harris (HHMI and JHU), Michael Häusser (UCL), Sonja Hofer (UCL), Edvard I Moser (NTNU), Josh Siegle (AI), Nicholas Steinmetz (UW), and Karel Svoboda (AI).

## Author contributions

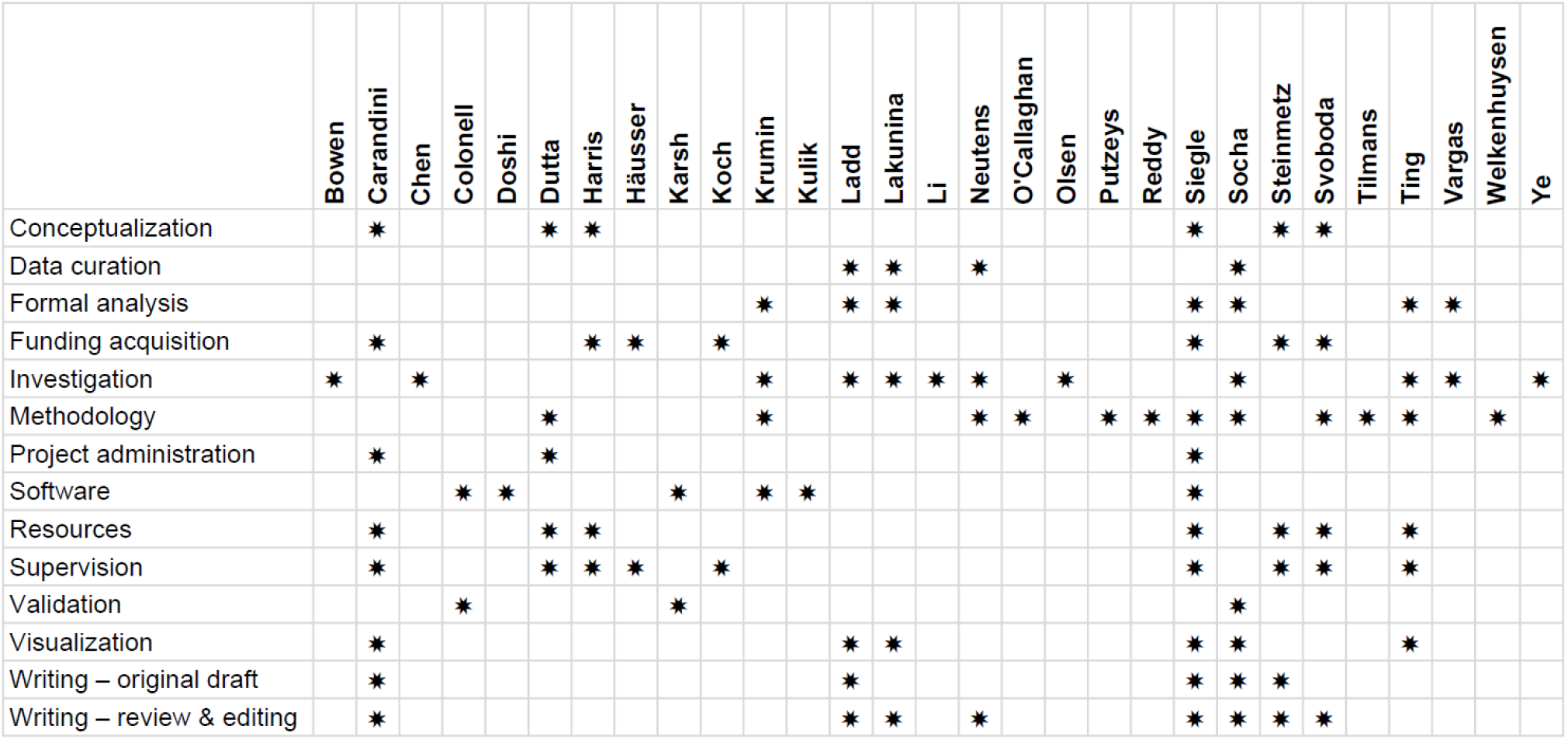

Author contributions according to the [CRediT](https://credit.niso.org/) taxonomy.

## Methods

### Design and fabrication

Neuropixels Opto probes involve a monolithic integration of a CMOS recording platform with a dual-color photonic platform. This design provides 384 readout channels for the 960 recording sites, along with two sets of 14 programmable light emitters^55^.

The foundation of the probe is the integrated circuit (IC) for electrophysiological recording, which is unchanged from the Neuropixels 1.0 probe and is built on a 130-nm SOI CMOS aluminum process with six metal layers. The probe outline and profile are defined with MEMS-like micromachining. The shank measures 10 mm in length, 70 µm in width, and 33 µm in thickness. The 12 x 12 µm titanium nitride (TiN) recording sites are arranged in a 2×480 linear array, with a pitch of 20 µm vertically and 48 µm horizontally.

The key innovation in the Neuropixels Opto probe is the integration of a photonic layer between the IC and the recording electrodes (**Suppl. Fig. S1)**. This was accomplished using plasma-enhanced chemical vapor deposition (PECVD) to create silicon nitride (SiN) waveguides. These waveguides are 150 nm thick, a dimension chosen to ensure single-mode operation at both 450 nm (blue) and 638 nm (red) wavelengths. We measured the propagation loss for these waveguides on 84 dies over a full 200 mm wafer, yielding 2.97 ± 0.10 dB/cm at 450 nm and 0.52 ± 0.01 dB/cm at 638 nm. To route the waveguides, circular bends with a 40 µm radius were used, with minimal losses of 0.006 ± 0.003 dB and 0.016 ± 0.001 dB per 90-degree bend for the two wavelengths.

To accommodate the photonics circuitry and the fiber block, we extended the original 5-mm probe base integrating the neural readout circuits with two wings of 3 mm for the fiber block area and 2 mm for the photonic switching area.

The fiber block area includes eight grating couplers, to which an 8-fiber block with a 127-µm pitch is aligned. The two central gratings couple the 450 nm and 638 nm light from the signal fibers, while the remaining six are used for active alignment and for measuring coupling losses. The average coupling loss at 450 nm was 5.5 dB, measured over 47 probes.

The photonic switching area includes two four-level, current-driven photonic binary switching trees, allowing the light routed to any of the 16 (2^4^) output waveguides (**Suppl. Fig. S2**). These switches are Mach-Zehnder interferometers that function as toggle switches, with thermal phase shifters that operate on the thermo-optic effect. For a 2π phase shift, their efficiency is 16 mW for 450 nm and 35 mW for 638 nm, and the switch has a bandwidth of 20 kHz. To select a specific emitter, four photonic switches must be driven, requiring a total of eight current sources for the two colors. The total insertion loss of the switching tree is 4.3 dB for the 450 nm wavelength and 3.5 dB for the 638 nm wavelength. After calibration, any of the 2×14 emitters can be selected.

The two sets of 14 waveguides are routed to the distal end of the shank and are spaced 100 µm apart, covering the first 1.5 mm from the tip of the shank. To achieve broad angular emission, we used higher-order, apodized Bragg gratings, which spread the emission over multiple diffraction peaks. This design choice precluded the use of a reflector, resulting in a coupling efficiency of 40%.

We expected the cumulative insertion losses from the fiber input to the emitter output to be ∼25 dB for the 450 nm wavelength and ∼16 dB for the 638 nm wavelength. The primary contributor to the high loss for blue light is the propagation loss. Measurements from 31 probes confirmed these expectations, with average total loss values of 26.5 dB for the 450 nm light and 17.0 dB for the 638 nm light.

To integrate the photonics circuitry while maintaining CMOS performance, we addressed two main challenges. First, the addition of the photonics, and in particular of the high-stress SiN waveguide layer, could cause the thin shank to bend excessively. To counteract this bend, we deposited a SiN compensation layer, and we used the SiN capping layer for fine-tuning, ultimately achieving a tip deflection < ±200 nm. Second, to prevent scattered light from the waveguides from interfering with the CMOS circuitry, we implemented a TiN/Al-based light-blocking layer. The oxide cladding layers of the PECVD SiN waveguides were kept as thin as possible to minimize their impact on shank bending and thickness.

### Light source

Except where indicated, the light source was a PXI-mounted laser module emitting light at 450 nm and 638nm (Quantifi Photonics, New Zealand), connected to the Neuropixels Opto probe via an optic fiber. However, the light source could in principle be replaced by any laser with the appropriate wavelength. For instance, as indicated in later sections, some experiments were performed with a separate laser from Oxxius (France).

After the electrical characterization measurements, the light power was calibrated based on the measured loss for each emitter, such that the output levels were consistent across emitters.

Emitter control was performed with SpikeGLX or with OpenEphys GUI, which were updated for the purpose. This functionality was integrated into the publicly available versions of these packages. Analog modulation of laser output was achieved by controlling a PXI-mounted National Instruments data acquisition module with a Python or Matlab script, with network messages used to synchronize light output with emitter selection.

### Electrical and optical characterization

The electrical and optical characterizations (**Fig. 2,Suppl. Fig. S3**) were performed at IMEC.

#### Electrical characterization

Measurements were performed in a grounded Faraday cage. The probe shank was immersed in phosphate-buffered saline (PBS) solution. The channels were configured to use external reference and x1000 gain. The external reference input was connected to the ground pad of the probe. First, we measured gain. We applied a sinusoidal test signal of 500 μV (peak to peak) at 1.5 kHz or 150 Hz (for AP or LFP band) to the PBS solution using a platinum (Pt) counter electrode. We recorded the probe signal and calculated the gain at the two frequencies. Second, we measured noise. We grounded the PBS solution via the external reference and ground contact pads on the flex cable. We recorded the probe signal and we calculated the integrated noise in the frequency bands for AP and LFP. We then divided the measured noise by the measured gain to obtain the input-referred noise.

#### Optical characterization

To measure the full probe loss, we inserted the shanks into an integrating sphere (AvaSphere-30-IRRAD, Avantes) and measured the output power, for a fixed input power, while sweeping from emitter 1 to emitter 14. To measure the emitter radiation pattern, we used a Nikon Eclipse microscope with a water immersion objective (Nikon Fluor, 60X/1.0NA), and a motorized 3D fiber coupling stage (based on the PI Q-545 linear stage). Images were acquired with a scientific CMOS camera (Hamamatsu Orca Flash) providing a field of view of 220 x 220 μm. To perform the measurement, the microscope field of view was centered on the emitter, and the objective was focused on the waveguide plane. A 200 μm z-scan was performed with a 4 μm step, and the imaged frames were saved into a 3D matrix.

These measurements were made on a test structure: a 1 cm long waveguide with a standard grating coupler on one side, and the emitter on the other side. Light was coupled into the structure with a horizontally placed 40 deg angle polished fiber, which was actively aligned to the standard grating coupler. The test structures were arranged in large blocks with many waveguides next to each other on a 25 μm pitch. For red light, we could not avoid coupling some light into the neighboring waveguides, resulting in small artifacts where a small amount of light was visible in neighboring emitters (**Fig. 2b**, *top*). These artifacts were specific to the test structure layout and the properties of the fiber coupling. They were not present at the emitters of fully constructed probes.

### Measurements in an optical phantom

The measurements of scattering in an optical phantom (**Suppl. Fig. S4**) were performed at University College London.

To measure the light projected by a Neuropixels Opto emitter onto individual planes (optical sectioning), we placed the probe at various distances from a fluorescent imaging plane, and imaged its fluorescence with a camera.

#### Fluorescent imaging plane

To construct the fluorescent imaging plane we coated a Superfrost Plus glass slide (Cat. No. 631-9483, Menzel, Avantor Sciences) with ∼10 µL of carboxyl Quantum Dots (Qdot 655 ITK, Cat. No. Q21321MP, Thermo Fisher). The Qdots were spread into a thin, even layer by gently placing a clean glass coverslip at the edge of the droplet and slowly dragging it across the surface to promote capillary action. We then placed the slide in a 45 °C incubator on a shaker for 2-3 minutes to promote adhesion of the Qdots to the slide surface and to allow the aqueous solvent to partially evaporate. A small drop (∼15 µL) of optical adhesive (NOA 81, Norland Products, Thorlabs) was subsequently applied over the dried Qdot layer. Using the same method as before, we used a coverslip to gently spread the adhesive into a thin, uniform film across the slide. The adhesive layer was then cured using UV light at 365 nm for ∼3 min to ensure complete crosslinking of the polymer. This curing step rendered the Qdot-coated slide water-tight and suitable for extended aqueous immersion. The final height of the coating was measured at ∼30 µm using a dial gauge with resolution of 10 µm.

#### Imaging

A micromanipulator (uMp-4, Sensapex) varied the distance between the probe and the imaging plane, and a camera (BFS-U3-50S5M-C. Teledyne FLIR) focused on the imaging plane using an air objective (UPlanFL N, 4x, NA 0.13, Olympus) acquired images through an optical filter (FF01-676/37-25, Semrock). The resulting resolution of the images was 2.3 µm/pixel. The probe was first immersed in water, to establish baseline measurements, and then in 1% milk. This medium was chosen because its estimated^62^ reduced scattering coefficient μ^′^_s_ (22.5 cm^-1^) is in the range of the values measured in rodent gray matter^62^ (20-30 cm^-1^).

### Spike sorting and quality metrics

In the recordings *in vivo* described below, spike sorting was performed with Kilosort^2,90^, typically accessed via SpikeInterface^91^. For quality control, we selected clusters that met these criteria: interspike-interval (ISI) violation ratio < 0.5; amplitude cutoff < 0.1; presence ratio > 0.8. These quantities are defined at https://spikeinterface.readthedocs.io/en/latest/modules/qualitymetrics.html.

### Activating local neural populations

The experiments demonstrating recording and activation of local neural populations (**Fig. 3,Suppl. Figs. S6-S10**) were performed at University College London. Experimental procedures were conducted according to the UK Animals Scientific Procedures Act (1986) under personal and project licenses released by the Home Office following appropriate ethics review.

#### Mice and viral strategy

The experiments described here were performed on 4 adult mice (aged 10-16 weeks at the time of headplate implantation): two males with wildtype background (C57BL/6, Charles River) and two females with double-transgenic background (Ai32^92^ x PV-Cre^93^, JAX #012569 and #008069). The 2 transgenic mice expressed the blue-sensitive opsin ChR2 in inhibitory (PV+) neurons. For the measurements presented here, 3 mice were injected with a virus expressing red-sensitive opsin ChRmine in CaMK2+ neurons^49^ (AAV-8-CaMKIIa-ChRmine-mScarlet-Kv2.1-WPRE, GVVC-AAV-194, Stanford Viral Core). An additional male with wildtype background (C57BL/6, Charles River) and no opsin expression was used as a control.

#### Main surgery

An initial surgery was performed to implant a headplate, perform a craniotomy, and inject a virus. Procedures were adapted from an established protocol^94^. Briefly, mice were injected with dexamethasone (i.m.) before surgery, Neuropixels Opto and then anesthetized with isoflurane (3% for induction; 1–1.5% for maintenance). Appropriate hydration and temperature control were provided. A steel headplate was attached to the skull and secured with dental cement (Super-Bond C&B, Sun Medical Co). The skin margins were attached to the cranium with tissue adhesive (Vetbond, 3M). A 3 mm craniotomy was performed, centered on the left primary visual area (VISp, ∼3.7 mm posterior and ∼3 mm lateral from bregma). A glass pipette (Drummond Scientific), beveled to form a ∼25-40 μm tip (EG-45 Microgrinder, Narishige), was lowered stereotaxically 150, 300, and 550 μm into the brain, to deliver 70 nL of viral vector solution (2×10^12^ vg/mL) at each depth (Nanoject II, Drummond Scientific), with 3 min pauses between depths, and a 5 min pause at the bottom. Injections were performed in 4-5 locations in VISp, 500-750 μm apart. The total volume of virus solution delivered was 840-1050 nL. The craniotomy was then covered with a removable window^95^ comprising two 3-mm circular cover glasses (#1) attached to a 5-mm circular cover glass (#1, Warner Instruments) using optical adhesive (Norland Optical Adhesive NOA 61, Thorlabs). The remaining exposed cranium and the skin margin were covered with cement (Super-Bond C&B). Post-operative treatment was provided for 3 days with carprofen in drinking water. Later, the mice were handled and habituated to the head-fixed recording rig for 30-60 min for at least 4 days before any recordings.

#### Widefield imaging

We waited 3–4 weeks for the virus to fully express and found the locations of virus expression using epi-fluorescence widefield imaging involving an illuminator (X-Cite DC200, Excelitas, Canada), a trinocular (Nikon C-TF), a 4x 0.13 NA air objective (UPlanFL N, Olympus), filter cubes for GFP and TurboFP635 (Chroma, VT), and a sCMOS camera (PCO.Edge 5.5 CLHS, Excelitas Canada Inc.). We then prepared a replacement glass window with holes in the appropriate locations.

#### Window replacement

To replace the glass window with one that had drilled holes At least 12 hours before the first recordings, we performed a brief procedure to replace the glass window. Mice were anesthetized using isoflurane (3% for induction; 1–1.5% for maintenance). The cement around the glass window was removed with a dental drill, and the new window was implanted in its place. We made an incision in the dura at the recording site to allow easier probe insertion, then covered the craniotomy with artificial dura^96^ (Duragel, Cambridge NeuroTech) and sealed the holes with Kwik-Cast (WPI, USA).

#### Recordings

A Neuropixels Opto probe with metal dovetail was mounted on a HHMI-designed probe holder and then on a 4-axis micromanipulator (uMp-4, Sensapex). In some recordings, we labeled the probe tip using Vybrant CM-DiI or DiO (V22888 or V22886, ThermoFisher). We lowered the probe to the brain surface and inserted it at 2 µm/s, typically reaching a depth of 1.2-1.4 mm. We then waited 15 min for the probe to settle, and started the recordings. During recordings, we controlled the probe and acquired signals using SpikeGLX, with standard gain settings of 500x and 250x for the AP and LFP bands. After recordings, we sealed the hole in the glass window with KwikCast, and we cleaned the probes by placing the shank in distilled water overnight. Occasionally, the probes required additional cleaning due to Duragel or tissue sticking to the shank; they were rinsed with a 1% Tergazyme solution for 30 min before being moved to distilled water.

#### Photostimulation

To activate neurons with light, we presented a 400 ms tapered square pulse of red light (638 nm). To minimize light artefacts^1^, the pulse had a smooth onset, ramping from 0 with half a cycle of a 40 Hz sine wave. The pulses were randomized across the 14 emitters, randomlyinterleaved with control trials (external illumination, visual stimulation, and gray screen), and repeated 40 times. The average inter-trial interval was 1.0 s. External optical activation was done using a 638 nm diode laser (LuxX 638-150, Omicron-Laserage Laserproducte, Germany). The laser light was delivered to the brain surface using a 200 mm, 0.22 NA patch cord (M122L02, Thorlabs), a collimator (F280FC-A, Thorlabs), and a focusing lens (f = 50mm, LA1213-A, Thorlabs) positioned ∼5 cm above the brain surface. Visual stimulation (full-field checkerboard, flickering at 4 Hz, 100% contrast, 1 s) was displayed on an LCD screen (LP097Qx1, LG).

#### Data processing

Signals in the AP band were filtered with a 300–12,000 Hz 3-pole Butterworth bandpass filter, followed by ADC phase-shift correction and common median subtraction. Sessions were spike-sorted with Kilosort4^90^ using default parameters. Sorting jobs were automatically generated and executed using custom Python packages adapted from https://github.com/spkware/spks. Quality metrics were calculated by adapting code from SpikeInterface^91^ and neurons were selected based on the criteria described in an earlier section. For multi-unit activity (MUA) we used less stringent criteria: amplitude cutoff < 0.1, presence ratio > 0.7, and ISI violations < 2.

#### Measurements of yield

To estimate the yield in terms of units per recording site we counted the number of neurons that passed quality control and were located in the visual cortex. We compared these numbers with those in the IBL database^68^ for recordings (performed with Neuropixels 1.0 probes) in the same general brain region (AP between -2200 and -3800, ML between -1800 and -3800, inserted at max 4100 deep), and we applied the same quality control metrics. The median yield for our experiments with Opto probes (0.23 units/site) resembled the median yield in IBL recordings (0.22 units/site).

#### Local field potentials

Neuropixels probes record local field potentials (LFPs) by filtering and digitizing voltage fluctuations synchronized with neural activity measurements, with the LFP band sampled at 2.5 kHz. LFPs were computed by bandpass filtering the raw signal (5–60 Hz) using non-causal, zero-phase delay filtering. Current Source Density (CSD) maps were generated from average LFPs using established methods^97^. LFPs from the uppermost and lowermost channels were duplicated, smoothed across adjacent channels with a weighted filter, and the second spatial derivative was calculated using a sampling interval of 40 µm. The resulting CSD data were linearly interpolated.

#### Width of activated region

To quantify the spatial spread of optogenetically evoked activity, we computed mean z-scored firing rates for each recording site and stimulating emitter, binned in 50 µm depth intervals. The resulting matrix was visualized as a heatmap, with axes representing depth relative to the probe tip and position of the stimulating emitter (**Fig. 3f**). We then calculated a single activation profile as a function of distance from stimulating emitter, by realigning relative to stimulation depth and averaging across all stimulation emitters located in cortex. We fitted this profile with a double-exponential decay function, and we computed the full width at half maximum (FWHM) from the fitted profile.

#### Histology

Following the recordings, the mice were perfused with 4% PFA (#28908, ThermoFisher). The brain was dissected and postfixed in PFA for 24 hours. Then, the brain was stored in 10% PBS for at least 48 hours before sectioning. We imaged full 3D stacks of the brains in a custom-made serial section two-photon tomography microscope^98^. Images were acquired using ScanImage (Vidrio Technologies) and the hardware was coordinated with BakingTray.

### Driving localized circuits

The spatially resolved neural inactivations (**Fig. 4** and **Suppl. Figs. S11-S13**) were performed at the University of Washington, in accordance with protocols approved by the Institutional Animal Care and Use Committee (IACUC).

#### Mice and viral strategy

The experiments described here were performed on 2 adult female mice and 1 adult male mouse (aged 19 weeks at the time of headplate implantation) with transgenic expression of GCAMP8s in CAMK2-positive neurons (CaMK2a-tTA.tetO-G8s). Mice were injected with a virus expressing red-sensitive opsin ChrimsonR-tdTomato under the control of the DLX2.0 enhancer^72-74^ (AAV-PHP.eB-DLX2.0-ChrimsonR-tdTomato, Addgene plasmid #229775). The virus (100 μL, 1.6×10^13^ vg/mL) was retro-orbitally injected in anesthetized (isoflurane 1-4% in O2) when the mice were 4–6 weeks old.

#### Implant surgery

Implant surgeries were performed after mice reached p48. Mice anesthetized with isoflurane (1-4% in O2) and subcutaneously administered analgesics carprofen (5 mg/kg) and lidocaine (2 mg/kg). The skin and periosteum were cleared to reveal the dorsal skull. The edges of the implant were sealed to the skull then secured to the skull with cyanoacrylate (VetBond, World Precision Instruments) to protect the underlying muscle. A 3-D printed recording chamber was implanted on top of the skull using dental cement (Metabond, Parkell). Fast-curing optical adhesive (Norland Optical Adhesive 81, Norland Products) was applied to the surface of the skull and cured with UV light. A titanium headpost (ProtoLabs) was then cemented to the posterior end of the recording chamber. Carprofen (0.05 mg/ml) was given for 2 days in water after surgery. Mice were allowed to recover in the home cage for at least one week before habituation and head-fixation.

#### Recordings

Over 3 hours before a recording session, the mouse was anesthetized (isoflurane 14% in O2) and a 2-3 mm craniotomy was performed over the left visual cortex and right motor cortex. Craniotomies were sealed with transparent Duragel (Dow Corning 3-4680 Silicone Gel). After recovery from anesthesia, mice were headfixed. The Neuropixels Opto probe was mounted on a micromanipulator and manually driven to the craniotomy at a 45° angle to accommodate an overhead laser. Real-time electrophysiological data was monitored as the probe was driven through into the brain. Insertions aimed to avoid blood vessels. If a probe could not successfully record from a craniotomy, a new surgery was performed at a later date. Once probes reached the brain’s surface, they were driven to their target depth at 200 μm/min. Probes were allowed to settle for 10 min at their final depth before recording data with SpikeGLX. We used internal tip referencing with gain settings of 1000x and 1000x for the AP and LFP bands. Probes were slowly removed from the brain (∼1 mm/min) at the end of recording. Probes were submerged in a 1% Tergazyme solution overnight and then rinsed in deionized water the next day to clean debris off the shank of the probe.

#### Photostimulation

To activate and inactivate neurons with light emitted from the probe, we presented a 250 ms tapered square pulse of red light (638 nm) from a randomly chosen emitter with an average inter-trial interval of 1.4 s. To minimize light artefacts^1^, the pulse was tapered, ramping linearly up for the first 25 and then down for the last 25 ms. Experiments in mice 2 and 3 were performed using an Oxxius LBX-638nm diode laser. Calibrations were performed to ensure both lasers supplied a similar amount of light to the probe during experiments.

#### Data processing

We used SpikeInterface^91^ to preprocess the raw data (decompression, phase shift, high-pass filter, and median subtraction). We then used Kilosort 4 (Ref. ^90^) for spike sorting. We calculated quality metrics, and selected neurons based on the criteria described in an earlier section.. We used Neuropyxels^99^ to plot raw waveforms from SpikeGLX data. Light-activated neuron-emitter pairs were defined by a >300% increase in firing rate during the 250 ms light stimulus compared to the pre-stimulus baseline with p< 0.05. Inactivated neuron-emitter pairs were identified as having at least a 50% reduction in firing rate during the stimulus period with p< 0.05. Modulation index was computed as (R1-R0)/(R1+R0) where R0 and R1 are mean firing rates before and during stimulus. When calculating modulation index (**Fig. 4e**), we averaged the mean firing rates (R0 and R1) for the two most proximal emitters superficial to each unit.

#### Histology

Following completion of recordings, mice were transcardially perfused with PBS (50 mL at 5 mL/min) followed by 4% paraformaldehyde in PBS. Brains were extracted and post-fixed in 4% PFA overnight at 4°C. Fixed brains were incubated in D OFF solution (5 mL DI water, 5 mL SHIELD-Buffer, 10 mL SHIELD-Epoxy) at 4°C. After 5 days of incubation in SHIELD OFF buffer, brains were transferred to SHIELD ON Buffer at 37°C for 24 hours. Samples were cleared in LifeCanvas Delipidation Buffer at 45°C with shaking (5 days for hemispheres), then washed overnight in PBS/0.02% sodium azide at 37°C. Cleared tissues were incubated in EasyIndex until transparent. Endogenous GCaMP8s and ChrimsonR-tdTomato were imaged via lightsheet microscopy. Samples were mounted in 2% agarose/EasyIndex, and z-stacks acquired through cortical regions to visualize calcium indicator and GABAergic neuron distributions. More information is available at https://lifecanvastech.com/beginners-guide-to-tissue-clearing-with-lifecanvas-products/.

### Optotagging nearby neurons

The subcortical optotagging experiments (**Figs. 5** and **6** and **Suppl. Figs. S14** and **S17**) were carried out at the Allen Institute, in accordance with protocols approved by the Institutional Animal Care and Use Committee (IACUC).

#### Mice and viral strategy

Experiments were performed on 26 adult mice (11 males, 15 females; aged 11-28 weeks at the time of headframe implantation). We used eight transgenic lines:

- Chat-IRES-Cre^100^ (JAX #031661)
- Chat-IRES-Cre-neo^100^ (JAX #006410)
- Sst-IRES-Cre (JAX #028864)
- Drd1a-Cre^101^ (JAX #037156)
- Adora2a-Cre (MMRRC #36158)
- Slc17a6-IRES-Cre^102^ (JAX #028863)
- Ntrk1-IRES-Cre (MMRRC #15500)
- Gad2-IRES-Cre^103^ (JAX #028867)

In addition, some mice received one or more of the following viruses injected stereotaxically at the titers listed below:

- pAAV-Syn-FLEX-rc(ChrimsonR-tdTomato) (Addgene #62723); 3.39e13 GC/mL
- pAAV-Ef1a-DIO-ChRmine-mScarlet-WPRE (Addgene #130998); 1.07e13 GC/mL
- pAAV-Ef1a-DIO-rsChRmine-oScarlet-Kv2.1-WPRE (Addgene #183529); 7.60e12 GC/mL
- hSyn-DIO-somBiPOLES-mCerulean (Addgene #154951); 2.41e12 GC/mL
- AiP14033: pAAV-AiE0779m_3xC2-minBG-CoChR-EGFP-WPRE3-BGHpA (Addgene #214852); 1.41e13 GC/mL
- AiP14035: pAAV-AiE0452h_3xC2-minBG-CoChR-EGFP-WPRE3-BGHpA (Addgene #214853); 5.15e13 GC/mL
- AiP14036: pAAV-AiE0743m_3xC2-minBG-CoChR-EGFP-WPRE3-BGHpA (Addgene #214854); 4.13e13 GC/mL

The example experiment (**Fig. 5**) involved Adora2a-Cre mice (MMRRC #36158) injected with viruses expressing CoChR-EGFP (Addgene #214852) and ChRmine-mScarlet (Addgene #130998).**Surgery**. Mice were anesthetized and placed in a stereotaxic frame. The dorsal scalp was removed, the skull leveled, and bregma located using tooling adapted from a previously described headframe and clamping system^104^. An outline of the implant location was etched using a custom tracing tool, and the assembled headframe was cemented in place. A craniotomy was performed using the traced implant shape as a guide, and the dura was removed. If the mouse was to receive viral injections, these were delivered stereotaxically through the craniotomy. Afterwards, the prepared 3D-printed SHIELD artificial skull^105^ was placed in the opening. The edges of the implant were sealed to the skull using a light curing cyanoacrylate adhesive (Loctite 4305) and reinforced with dental cement. Finally, a removable plastic cap was placed over the well to protect the implant’s silicone coating. After at least one week of recovery, and prior to the first recording, the mouse was anesthetized and placed in a stereotaxic frame. The layer of durable silicone covering the SHIELD implant was removed. A ground wire was inserted into the grounding hole in the implant and pushed forward until it rested on the surface of the brain. A Duragel mixture was then prepared and poured over the implant to a thickness of at least 1 mm, then allowed to cure for at least 24 hours^106^.

#### Recordings

To allow post-hoc identification of probe tracks, probes were coated with DS-DiD (2mM in ethanol; ThermoFisher Product #D12730) by immersing them in a well filled with dye. Each probe was dipped five times to ensure adequate coating. Each Neuropixels Opto probe was mounted on a 3-axis micromanipulator (New Scale Technologies, Victor, NY) on a custom modular insertion system. We used Pinpoint^107^ to select the appropriate insertion coordinates and approach angle for the desired target structure. Probes were manually driven to the appropriate hole in the SHIELD implant and slowly lowered toward the surface of the brain. The operator observed real-time continuous electrophysiological signals to identify when the probe entered the brain. If the probe needed adjustment when attempting to insert (e.g. to avoid blood vessels), the probe was completely retracted out of the Duragel to prevent probe bending. If a probe could not be inserted into its assigned hole, another target hole was selected. Once all probes reached the brain surface, each probe was automatically inserted to its target depth at a speed of 200 μm/min. Once all probes reached their final depth, they were allowed to settle for 10 minutes, and photo documentation of the inserted probes was captured. Neuropixels data was acquired using the Open Ephys GUI^57^ with gain settings of 500x and 250x for the AP and LFP bands. Videos of the eye, face, and/or body were acquired with USB3 cameras (Teledyne FLIR) and Bonsai software^108^. At the end of the recording session, probes were slowly retracted from the brain (∼1 mm/min). To remove debris, probes were submerged in a 1% Tergazyme solution overnight then rinsed in deionized water the next day.

#### Optotagging

To identify recorded units reliably activated by light, we presented 20 Hz trains of 10 ms laser pulses with an average inter-trial interval of 300 ms. Blue light (450 or 473 nm) or red light (638 nm) was delivered via the PXI-mounted laser module or a two-channel laser combiner (Oxxius, France). Optotagged units were identified by their significant increase in firing rate to at least 4 pulses from at least one emitter (p < 0.05, Holm-Sidak adjustment for multiple comparisons), a maximum spike latency of 8 ms (defined as the time when the spike rate reached two standard deviations above baseline), and a mean response reliability > 0.3 (proportion of trials with at least one spike evoked during the laser presentation). If units were tagged by both blue and red light, they were considered red-tagged: red-shifted opsins can be activated by blue light, but blue-shifted opsins are insensitive to red light.

#### Data processing

Neuropixels raw data files were compressed using WavPack^109^ and transferred to an Amazon S3 bucket. We used a custom Nextflow pipeline running on the Code Ocean compute platform to run preprocessing, spike sorting, and curation. First, the data was decompressed and denoised using phase shift, highpass filter, and median subtraction. Then, spike sorting was performed using Kilosort 2.5 (Ref. ^2^). Units with waveforms unlikely to originate from action potentials were automatically labeled as “noise” using pre-trained models from the UnitRefine toolbox. Finally, we calculated quality metric for each unit and applied the quality control criteria mentioned in an earlier section. Each unit’s location along the shank was estimated based on monopolar triangulation using the waveform amplitudes for recording sites within 75 μm of the peak. All these calculations were made with SpikeInterface^91^.

### *In vitro* characterization

Experiments to characterize enhancer virus expression patterns (**Suppl. Fig. S15**) and evoked photocurrents (**Suppl. Fig. S16**) were carried out at the Allen Institute in accordance with protocols approved by the Institutional Animal Care and Use Committee.

#### AAV vector construction

AiP14033 (Addgene plasmid #214852), AiP14035 (Addgene plasmid #214853), and AiP14036 (Addgene plasmid #214854) were created by subcloning the CoChR-EGFP fragment from Addgene plasmid #59070 (a gift from Dr. Edward Boyden) using restriction enzyme digestion (BamHI/EcoRI) and ligation into AiP13044 (Addgene plasmid #191706), AiP12982 (Addgene plasmid #191709), and AiP13038 (Addgene plasmid #191720). AiP13276 (Addgene plasmid #214842) was used as a control vector for comparing enhancer driven ChR2(H134R)-EYFP vs enhancer driven CoChR-EGFP. Plasmid integrity was verified via Sanger DNA sequencing. All AAV plasmids were propagated in NEB stable E. coli at 30°C growth condition to prevent spurious DNA rearrangements.

#### AAV packaging and titer determination

Smallscale crude AAV preps were generated using the HEK293 cell culture and triple transfection method as described elsewhere^110^. Viral preps were purified by iodixanol gradient centrifugation and subjected to ddPCR for titer determination. Stereotactic injection of AAV vectors

#### Stereotaxic injection of AAV vector

For stereotaxic injection surgery, adult C57Bl/6J mice were deeply anesthetized to a surgical plane using an isoflurane vaporizer and placed into the stereotaxic injection frame. AAV virus was injected bilaterally into the dorsal striatum region (dSTR) using the following coordinates (in mm) relative to bregma: AP = 0.8, ML = 2.2-2.3 (on either side), and DV = 3.0-3.2. A total volume of 500 nL (titer of 1e9-2e9 GC/mL) was delivered at a rate of 50 nL per pulse with a Nanoject II pressure injection system. Before incision, the animal was injected with Bupivacaine (2-6 mg/kg) and post injection, the animal was injected with ketofen (2-5 mg/kg) and Lactated Ringer’s Solution; LRS (up to 1 mL) to provide analgesia. Adult mice that underwent in vivo virus injections were euthanized at 1-2 months post injection and transcardially perfused with 1XPBS followed by 4% paraformaldehyde (PFA); the brains were dissected for further analysis.

#### Tissue processing for slide-based epifluorescence imaging

Mice were anaesthetized with isoflurane and perfused transcardially with 10 mL of 0.9% saline, followed by 50 mL of 4% PFA. The brain was removed, bisected along the midsagittal plane, placed in 4% PFA overnight and subsequently moved to a 30% sucrose solution until sectioning. 30 μm sections were obtained using a freezing, sliding microtome. Mid-sagittal sections were collected and stained with DAPI ((4’,6-diamidino-2-phenylindole dihydrochloride) and/or propidium iodide (PI) to label nuclei and to reveal cellular profiles, respectively. Stained tissue sections were slide mounted using Vectashield hardset mounting medium (Vector Laboratories, catalog # H-1400-10) and allowed to dry for 24 hours protected from light. Once the mounting medium hardened, the slides were scanned with Aperio VERSA Brightfield epifluorescence microscope (Leica) in the UV, green, and red channels, illuminated with a metal halide lamp.

#### Immunohistochemistry

Brain slices were fixed in 4% PFA in phosphate buffered saline (PBS) at 4 °C overnight or up to 48 hours and then transferred to 1XPBS with 0.01% sodium azide as a preservative. Fixed slices were thoroughly washed with PBS to remove residual fixative, then blocked for 1 hr at room temperature in 1XPBS containing 5% normal goat serum and 0.2% Triton-X 100. After blocking, slices were incubated overnight at 4 °C in blocking buffer containing primary antibodies chicken IgY anti-GFP (Aves, 1:2000) and mouse anti-ChAT IgG1 for cholinergic cells (Atlas Labs, 1:1000). Following the overnight incubation, slices were washed for 15 min three times with 1XPBS and then incubated for 1 hr at room temperature in dye-conjugated secondary antibodies (1:1000) including Alexa Fluor 488 (goat antichicken), Alexa Fluor 647 (goat anti-mouse). Slices were washed for 15 min three times with 1XPBS, followed by 5ug/mL DAPI nuclear staining for 15 min. The slices were then dried on glass microscope slides and mounted with FluoromountG (SouthernBiotech, Birmingham, AL). Slides were stored at room temperature in the dark prior to imaging. Whole slice montage images were acquired with NIS-Elements imaging software on a Nikon Eclipse Ti2 Inverted Microscope System equipped with a motorized stage and epifluorescence illumination with standard DAPI, FITC, TRITC, Cy3, and Cy5 excitation/emission filter cubes.

Specificity of enhancer activity for the target striatal cell subclass or type was quantified as reporter and marker Ab double positive neuron count divided by total reporter Ab positive neuron count and multiplied by 100 to obtain a percentage. The main ROI for analysis was in the center of the dorsal striatum region. A minimum of 100 neurons were counted in the ROI. Completeness of labeling was quantified as reporter and marker Ab double positive neuron count divided by total marker positive neuron count and multiplied by 100 to obtain a percentage.

#### *In vitro* electrophysiology and optogenetic stimulation

Enhancer AAV injected mice were deeply anaesthetized by intraperitoneal administration of Avertin (20 mg kg^−1^) and were perfused through the heart with carbogenated (95% O_2_/5% CO_2_) artificial aCSF consisting of (in mM): 92 N-methyl-D-glucamine (NMDG), 2.5 KCl, 1.25 NaH_2_PO_4_, 30 NaHCO_3_, 20 4-(2-hydroxyethyl)-1-piperazineethanesulfonic acid (HEPES), 25 glucose, 2 thiourea, 5 Na-ascorbate, 3 Na-pyruvate, 0.5 CaCl_2_·4H_2_O and 10 MgSO_4_·7H_2_O. Brains were sliced at 300μm thickness on a vibratome (VT1200S, Leica Biosystems or Compresstome VF-300, Precisionary Instruments) using a zirconium ceramic blade and following the NMDG protective recovery method^111^. Mouse brains were sectioned in the coronal plane such that the angle of slicing was perpendicular to the pial surface. After sections were obtained, slices were transferred to a warmed (32–34 °C) initial recovery chamber filled with NMDG aCSF under constant carbogenation. After 12 min, slices were transferred to a chamber containing HEPES holding aCSF solution consisting of (in mM): 92 NaCl, 2.5 KCl, 1.25 NaH2PO4, 30 NaHCO3, 20 HEPES, 25 glucose, 2 thiourea, 5 sodium ascorbate, 3 sodium pyruvate, 2 CaCl2·4H2O and 2 MgSO4·7H2O, continuously bubbled with 95% O2/5% CO2. Slices were held in this chamber until use in acute patch clamp recordings or until fixed in 4% PFA for later histological processing. Mouse brains were sectioned in the coronal plane such that the angle of slicing was perpendicular to the pial surface. After sections were obtained, slices were transferred to a warmed (32–34 °C) initial recovery chamber filled with NMDG aCSF under constant carbogenation. After 12 min, slices were transferred to a chamber containing HEPES holding aCSF solution consisting of (in mM): 92 NaCl, 2.5 KCl, 1.25 NaH2PO4, 30 NaHCO3, 20 HEPES, 25 glucose, 2 thiourea, 5 sodium ascorbate, 3 sodium pyruvate, 2 CaCl2·4H2O and 2 MgSO4·7H2O, continuously bubbled with 95% O2/5% CO2. Slices were held in this chamber until use in acute patch clamp recordings or until fixed in 4% PFA for later histological processing.

Brain slices were placed in a submerged, heated (32-34°C) chamber that was continuously perfused with fresh, carbogenated aCSF consisting of (in mM): 119 NaCl, 2.5 KCl, 1.25 NaH_2_PO_4_, 24 NaHCO_3_, 12.5 glucose, 2 CaCl_2_·4H_2_O and 2 MgSO_4_·7H_2_O, 1 kynurenic acid, 0.1 picrotoxin (pH 7.3-7.4). Neurons were visualized with an upright microscope (Scientifica) equipped with infrared differential interference contrast (IR-DIC) optics. Glass patch clamp pipettes were pulled to an open tip resistance of 2-6 MΩ when filled with the internal recording solution consisting of (in mM): 110.0 K-gluconate, 10.0 HEPES, 0.2 EGTA, 4 KCl, 0.3 Na_2_-GTP, 10 phosphocreatine disodium salt hydrate, 1 Mg-ATP, 20 mg/mL glycogen, 0.5U/mL RNase inhibitor (Takara, 2313A), 0.5% biocytin and 0.02 Alexa 594 or 488 – pH adjusted to 7.3 with KOH. Whole cell somatic recordings were acquired using a Multiclamp 700B amplifier and custom acquisition software written in Igor Pro (MIES, https://github.com/AllenInstitute/MIES). Electrical signals were digitized at 50 kHz by an ITC-18 (HEKA) and were filtered at 10 kHz. The pipette capacitance was compensated, and the bridge was balanced during the current clamp recordings.

To measure peak photocurrent amplitude of ChR2(H134R)-EYFP vs CoChR-EGFP-expressing striatal neuron types, the area with maximal native reporter fluorescence was first identified. As labeling appeared as dense fluorescent neuropil rather than clearly discernable individual MSN somata, neurons with healthy-appearance under IR-DIC within this densely fluorescent region were selected for voltage clamp recordings with a holding command set to -70 mV (or -60 mV in the case of striatal cholinergic interneurons). Light was delivered from a mercury arc lamp attached to light guide directed through the 40X 0.8 NA water-immersion microscope objective equipped with a blue excitation spectrum bandpass filter cube. The power density was set to elicit maximal (saturating) photocurrent amplitude (50 mW/mm^2^). Recordings in the current clamp mode were obtained for select neurons to measure frequency– current curves to support striatal neuron type identity and signature firing properties.

## Data and code availability

The data and code used for this study will be made available in an open repository before publication.

## Supplementary Figures

**Figure S1.**
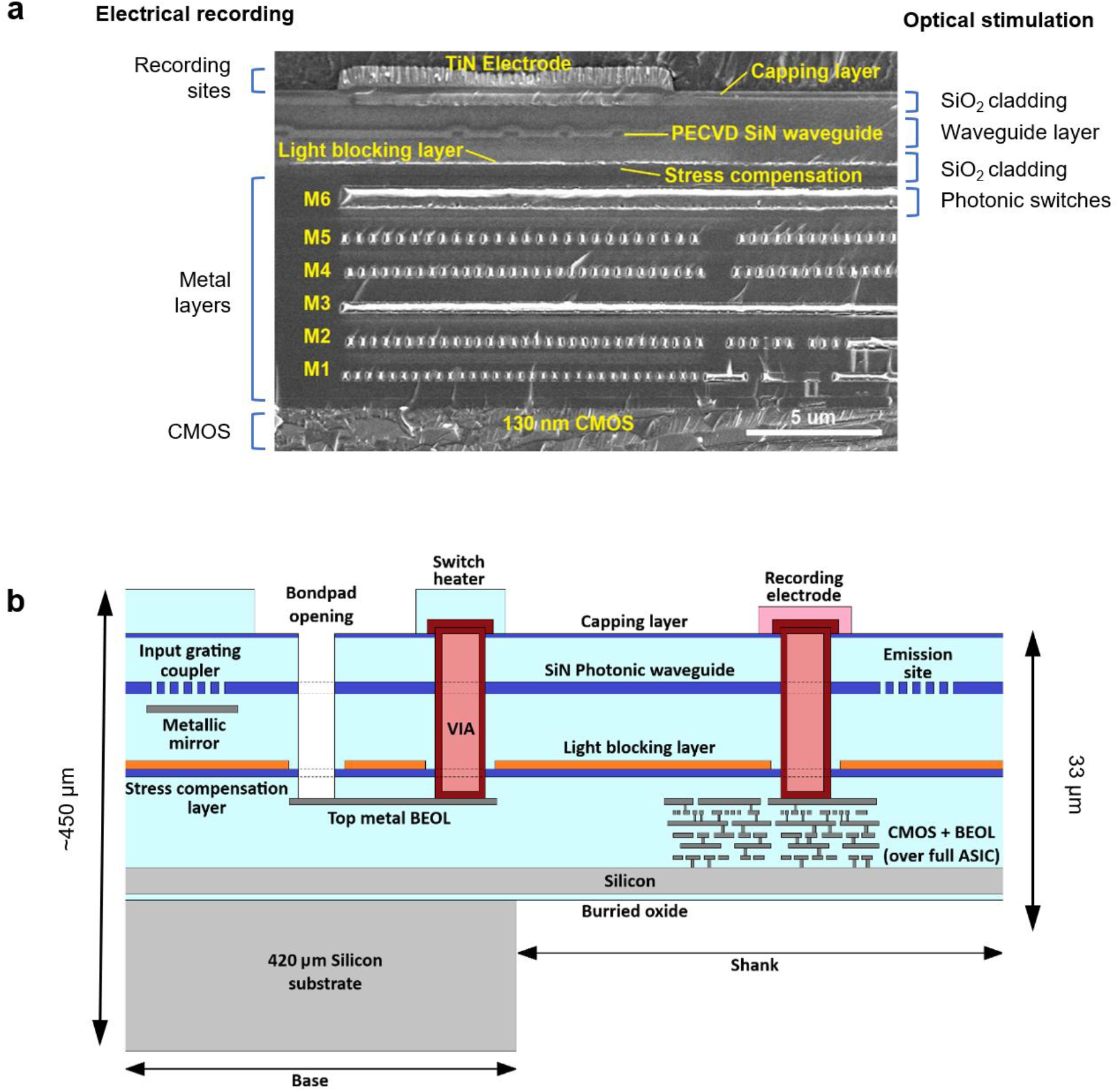
The integrated CMOS/Photonics platform. **a**. Cross-section of the probe shank obtained with a scanning electron microscope (SEM) showing the monolithically integrated platform combining the CMOS platform (for electrical recording) and Si_x_N_y_-based photonics (for optical stimulation). *Left labels*: layers used for the recording sites and readout IC, based on the 130-nm SOI CMOS Aluminum back-end-of-line process (Al BEOL) that was developed for the Neuropixels 1.0 probe. They point to the 130-nm CMOS layer (*bottom*), the 6 Aluminum metal layers (*middle*) and the Titanium nitride electrode (*top*). *Right labels*: the photonic modules that were added between the M6 metal layer of the CMOS BEOL and the recording electrode to enable optical stimulation. The photonics also use the M6 metal layer for controlling the photonic switches that are used to select the optical emitter. **b**. Schematic of the probe build-up, illustrating the photonic modules that were added to the CMOS platform, which include: grating couplers handling the light coupling from a fiber to the chip; an underlying reflector layer (metallic mirror); a Si_x_N_y_ photonic waveguide module; a full TiN/Al/TiN light blocking layer that is only interrupted for the VIA and bond pads, to protect the CMOS circuits from laser light scattered by the photonic waveguides; a heater module to make thermo-optical switches; a stress compensation layer to cancel the stress induced by the extra photonics layers, keeping the shank tip deflection within the spec of ±200 μm; and a deeper VIA etch to contact the heaters and electrodes.

**Figure S2.**
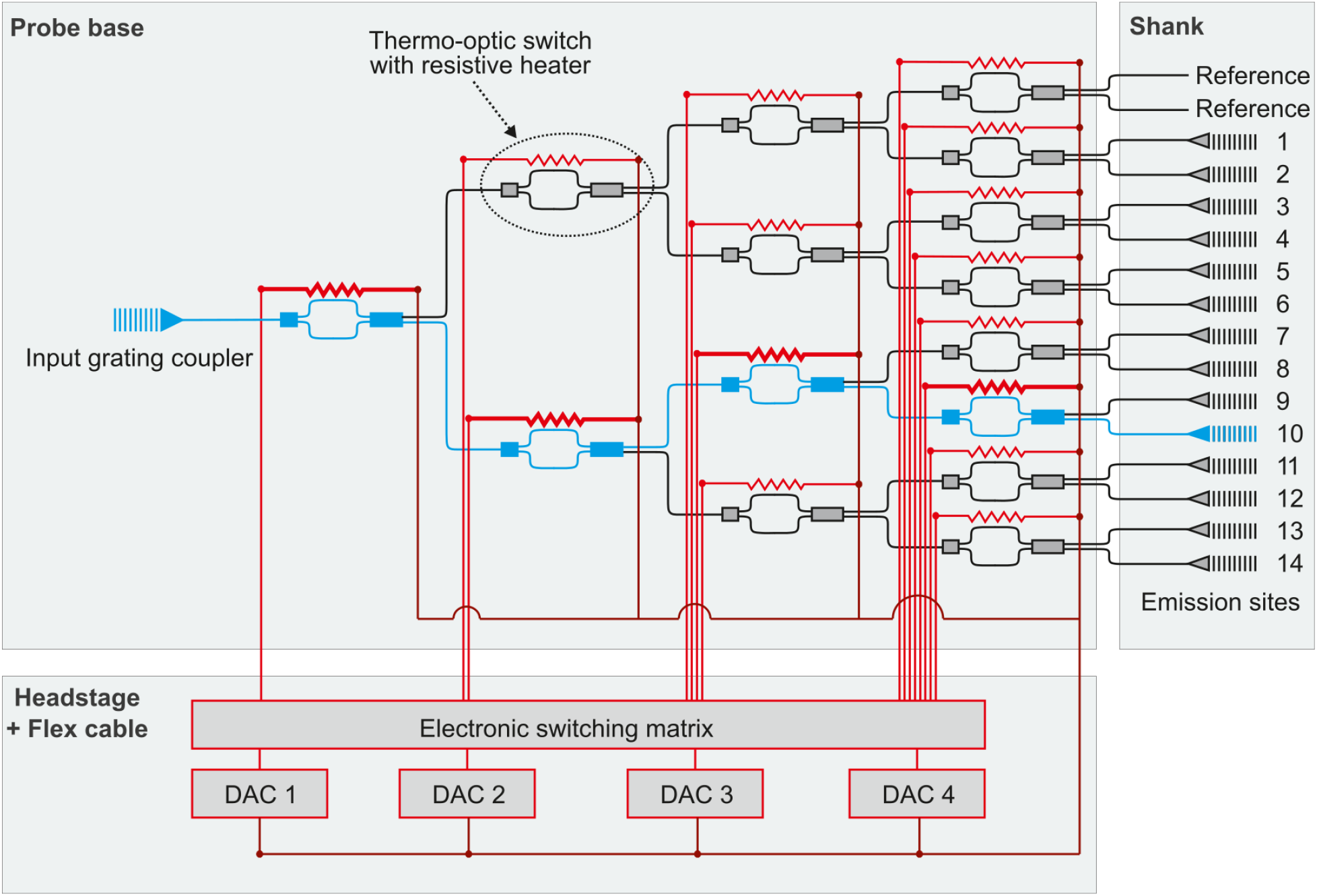
Switching tree. Schematic of switching tree architecture, which distributes light from a single input grating coupler to one of fourteen waveguides on the shank. A network of 15 thermo-optic switches is controlled by four digital-to-analog converters (DACs), allowing light to be routed to each emitter, or to reference sites used for calibration. Each probe contains a separate switching tree for blue and red light.

**Figure S3.**
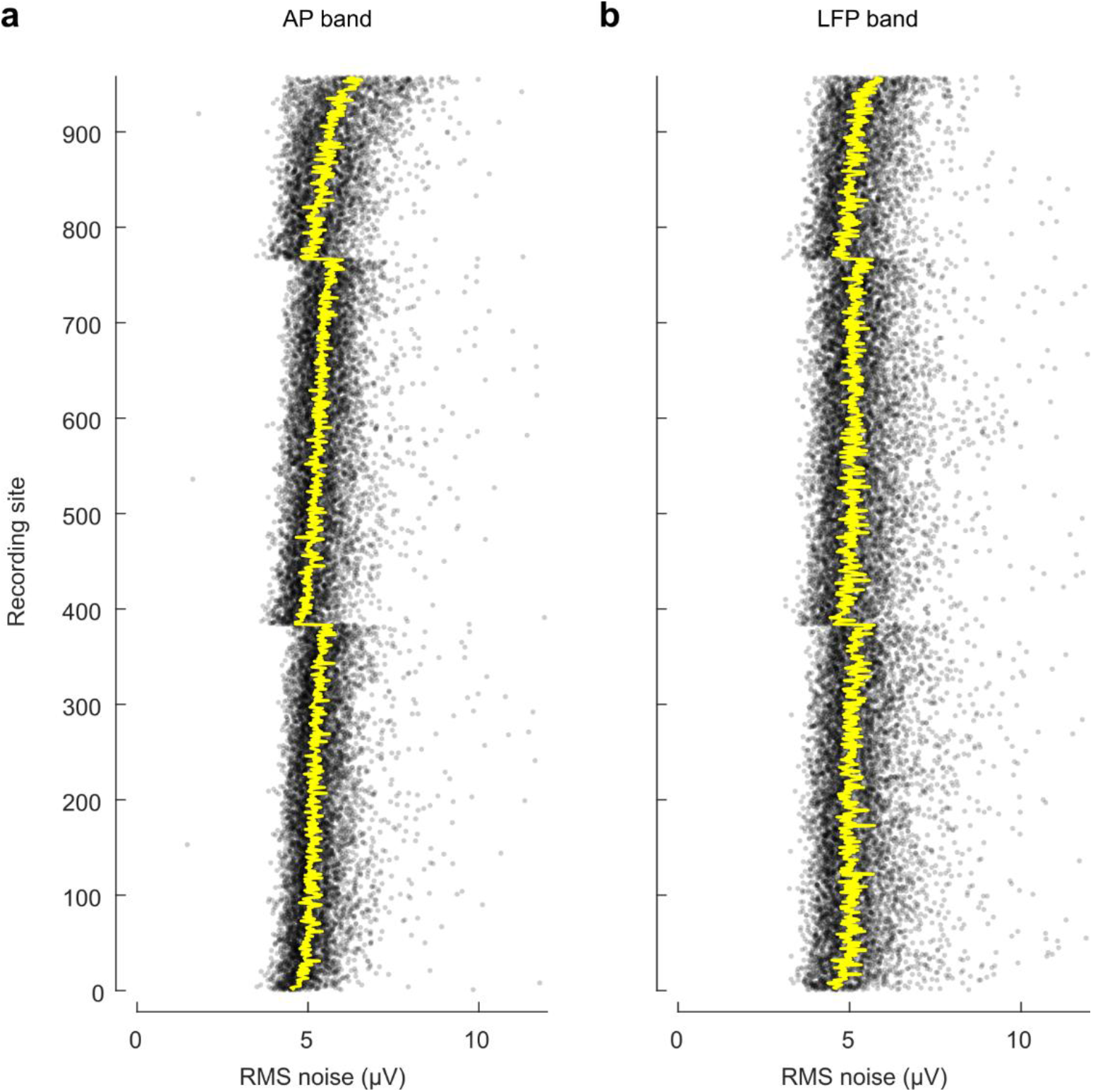
Electrical characterization. **a**. Input-referred RMS noise levels, measured in saline for 957 recording sites from *N* = 21 probes, in the AP band. Each dot represents a measurement for one site for one probe. Yellow line indicates the median across all probes. **b**. Same, for the LFP band.

**Figure S4.**
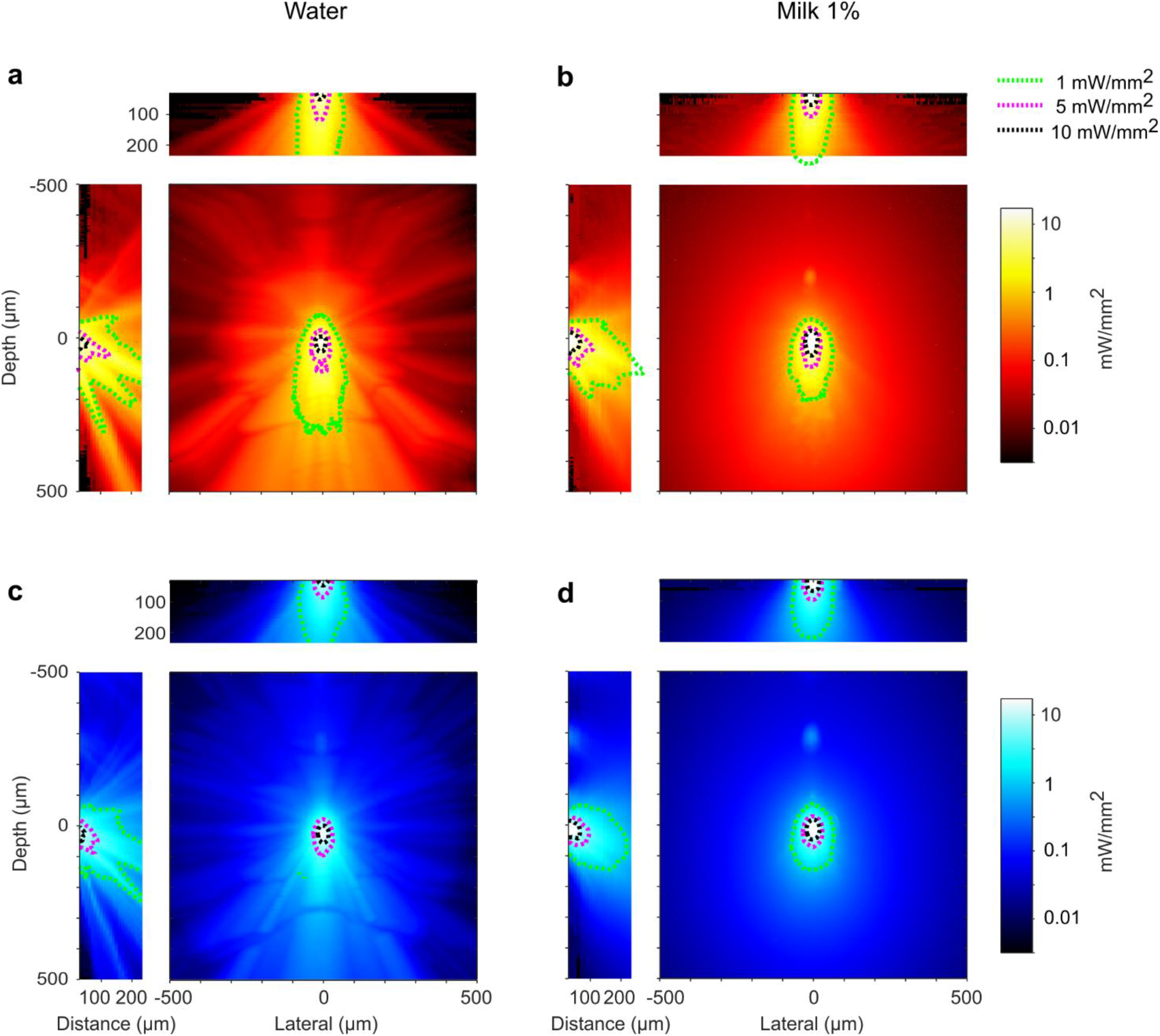
Light scattering in an optical phantom. To measure the 3D distribution of light projected by a Neuropixels Opto emitter, we constructed a fluorescent imaging plane (optical sectioning), and we sequentially varied the distance between the plane and the probe. **a**. Maximal intensity projections of the reconstructed volume for a red emitter, when the probe was immersed in water. Contours show intensity levels at 1, 5, and 10 mW/mm^2^. **b**. Same, when the probe was immersed in an optical phantom (1% milk) whose estimated^62^ reduced scattering coefficient μ^′^_s_ (23 cm^-1^) is in the range measured in rodent gray matter^62^ (20-30 cm^-1^). **c**,**d**. Same as a,b, for a blue emitter.

**Figure S5.**
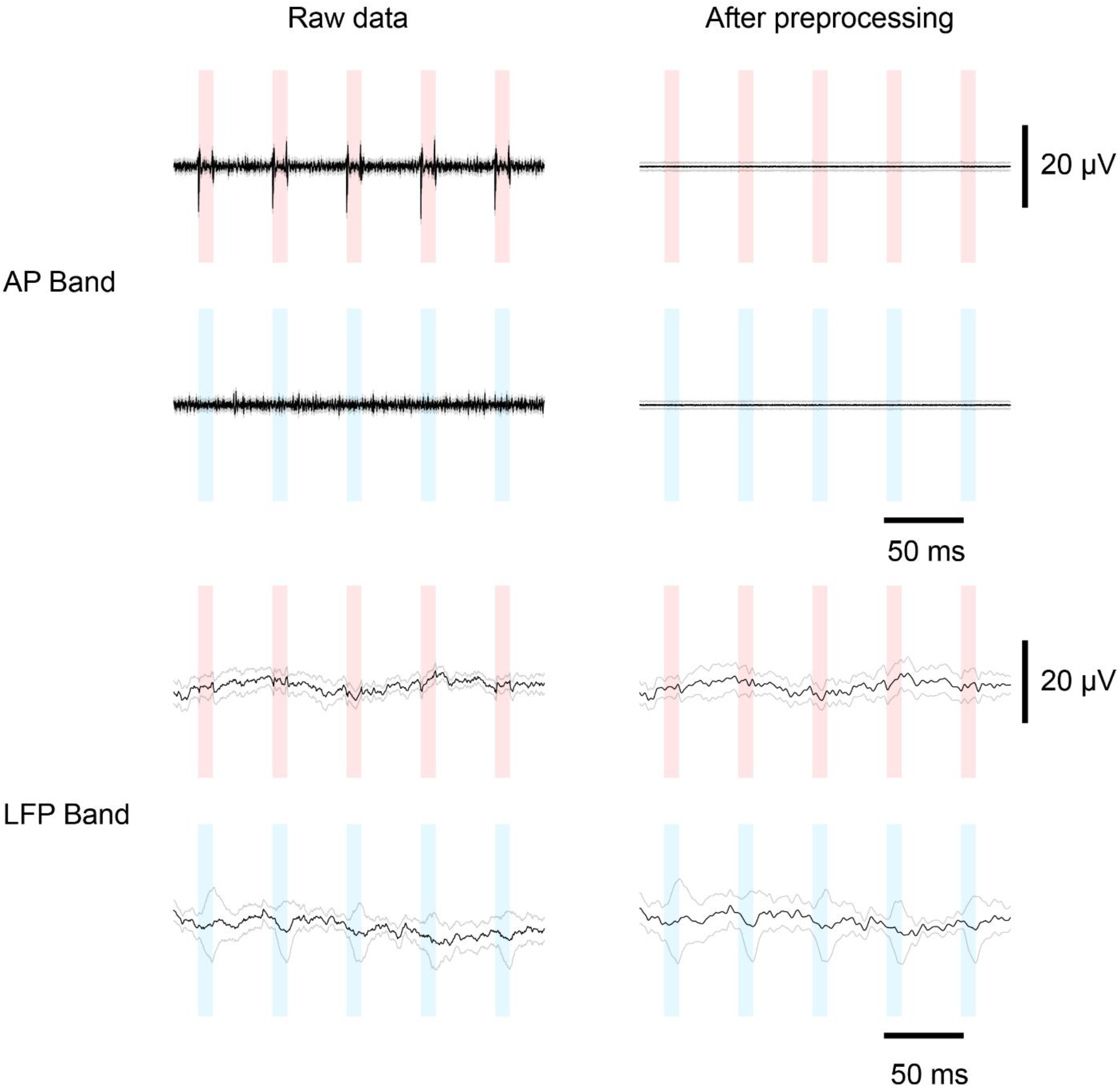
Measurement and removal of light artifacts. Measurements performed in brain tissue, showing light artifact for 10 ms red and blue light pulses at 100 μW, before and after standard pre-processing steps (average of 30 trials, 384 recording sites). The red and blue bars indicate periods of light activation (20 Hz train of 10 ms light pulses). The artifact seen for red pulses is highly uniform across recording sites, allowing it to be completely removed by phase shifting and median subtraction of the AP band data (see Methods for details). No such artifact was seen for blue light. In the LFP band, the small transients at the onset and offset of each pulse of red light are removed by applying a bandpass filter between 0.1 and 300 Hz.

**Figure S6.**
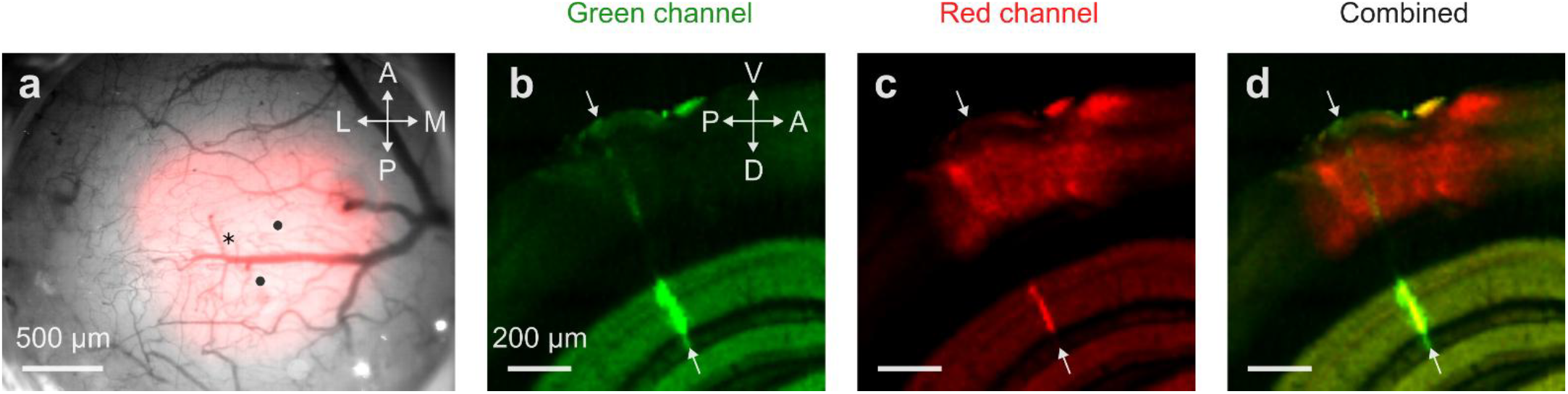
Opsin expression and probe track in the cerebral cortex. **a**. Widefield image of the visual cortex in Mouse I (see **Fig. 3**), showing the pattern of blood vessels (*gray*), the expression of ChRmine-mScarlet (*red*) and the location of the 3 probe insertions (*black symbols*). The *asterisk* marks the probe insertion shown in the subsequent panels. **b**. Example coronal section obtained with serial section two-photon tomography after perfusion. This image shows the green channel, revealing the track of the Neuropixels Opto probe (*arrows*), which had been immersed in DiO before insertion. The background expression in the hippocampus is due to autofluorescence. **c**. The same coronal section, viewed in the red channel, reveals expression of ChRmine-mScarlet in a patch of cerebral cortex but not in the underlying hippocampus. The DiO marking the probe track results in a faint additional red signal, particularly visible in the hippocampus (*arrows*). **d**. Superposition of the previous two images, showing the probe track in green and opsin expression in red.

**Figure S7.**
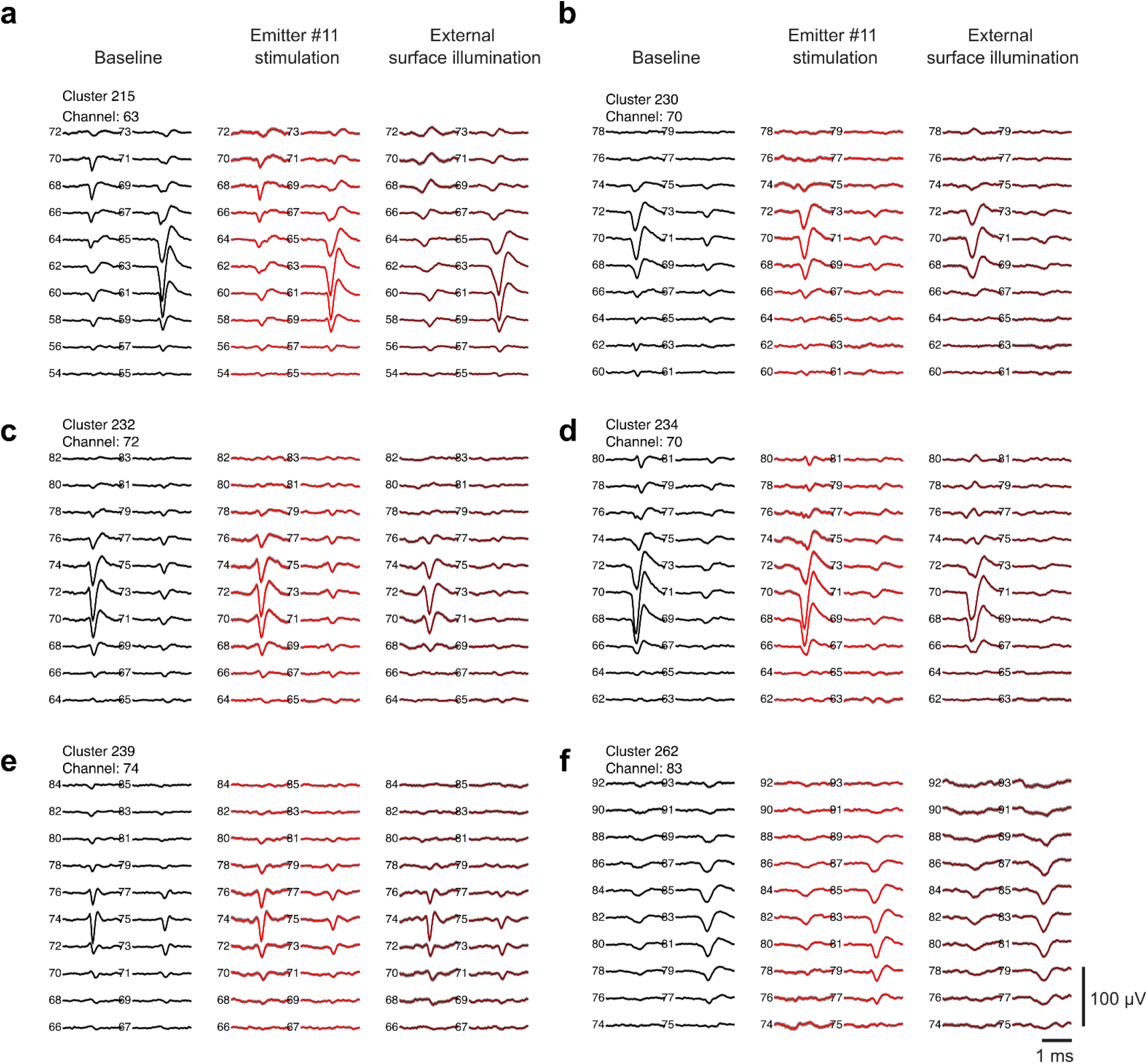
Spike shape invariance under light stimulation. Example spike waveforms from six clusters that passed quality control. These clusters appear in **Fig. 3b,c** and are stimulated by the same emitter (#11). For each cluster, the three panels show the spike shape recorded at 10 x 2 sites at baseline (1 s before emitter onset, *left*), during 400 ms emitter stimulation (*middle*), and during 400 ms external surface illumination (*right*). Each trace is the average of 100 spike waveforms. Shaded area (hardly visible) shows ± 1 s.e.

**Figure S8.**
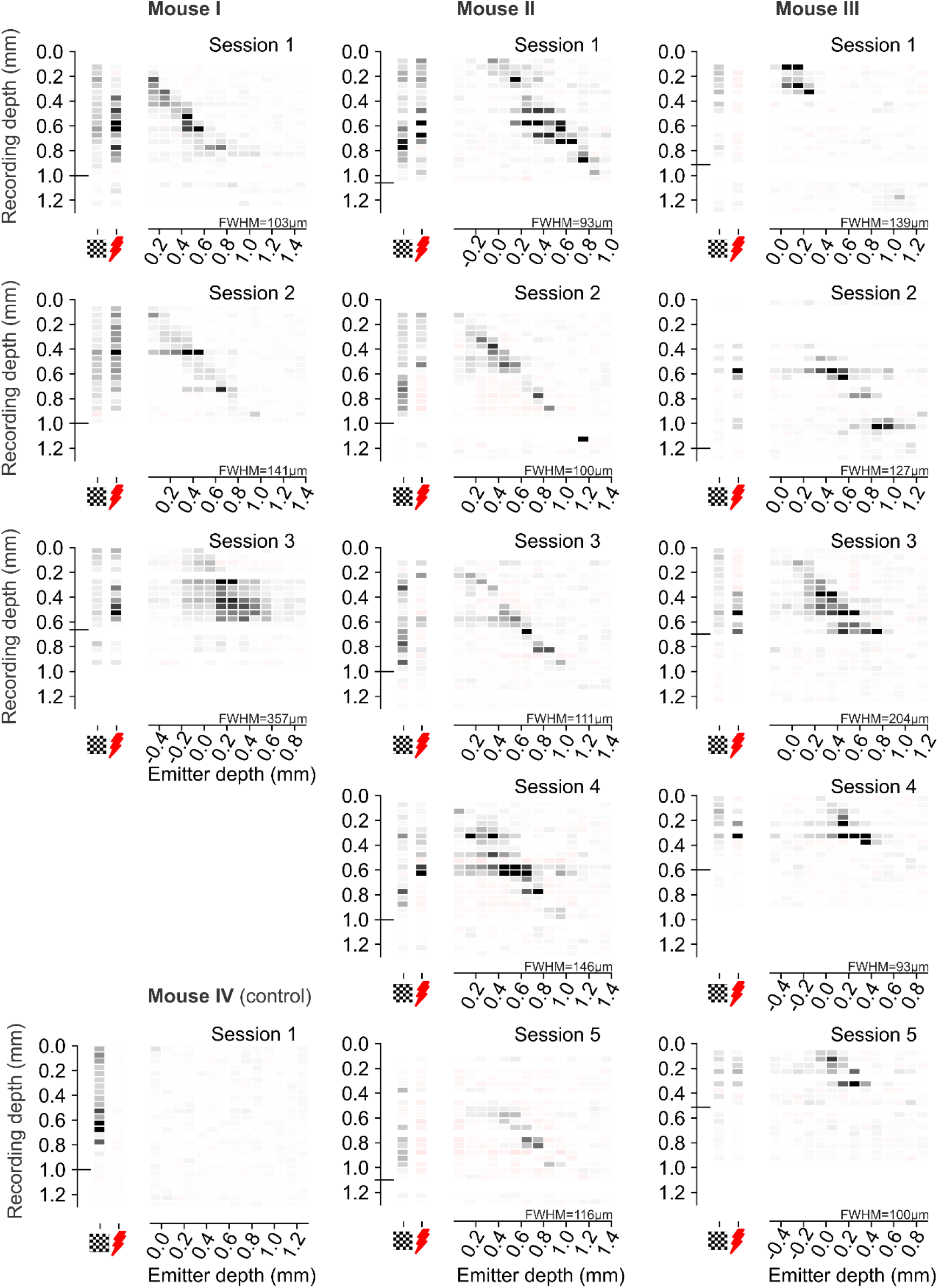
Additional examples of local circuit activation. Results of all the sessions with the experiments shown in **Fig. 3**. Format is as in **Fig. 3f**, showing average over time of response during stimulation with visual stimulus, surface laser, and single emitters (*abscissa*), at different cortical depths (*ordinate*). The horizontal line indicates the putative cortical border. In different insertions the top emitter was at different cortical depths, resulting in different ranges for the abscissa. Mouse IV is a control mouse that did not express any opsin. Accordingly, light stimuli (external or via the emitters) did not elicit any activity. More details on this control are in **Suppl. Fig. S10**.

**Figure S9.**
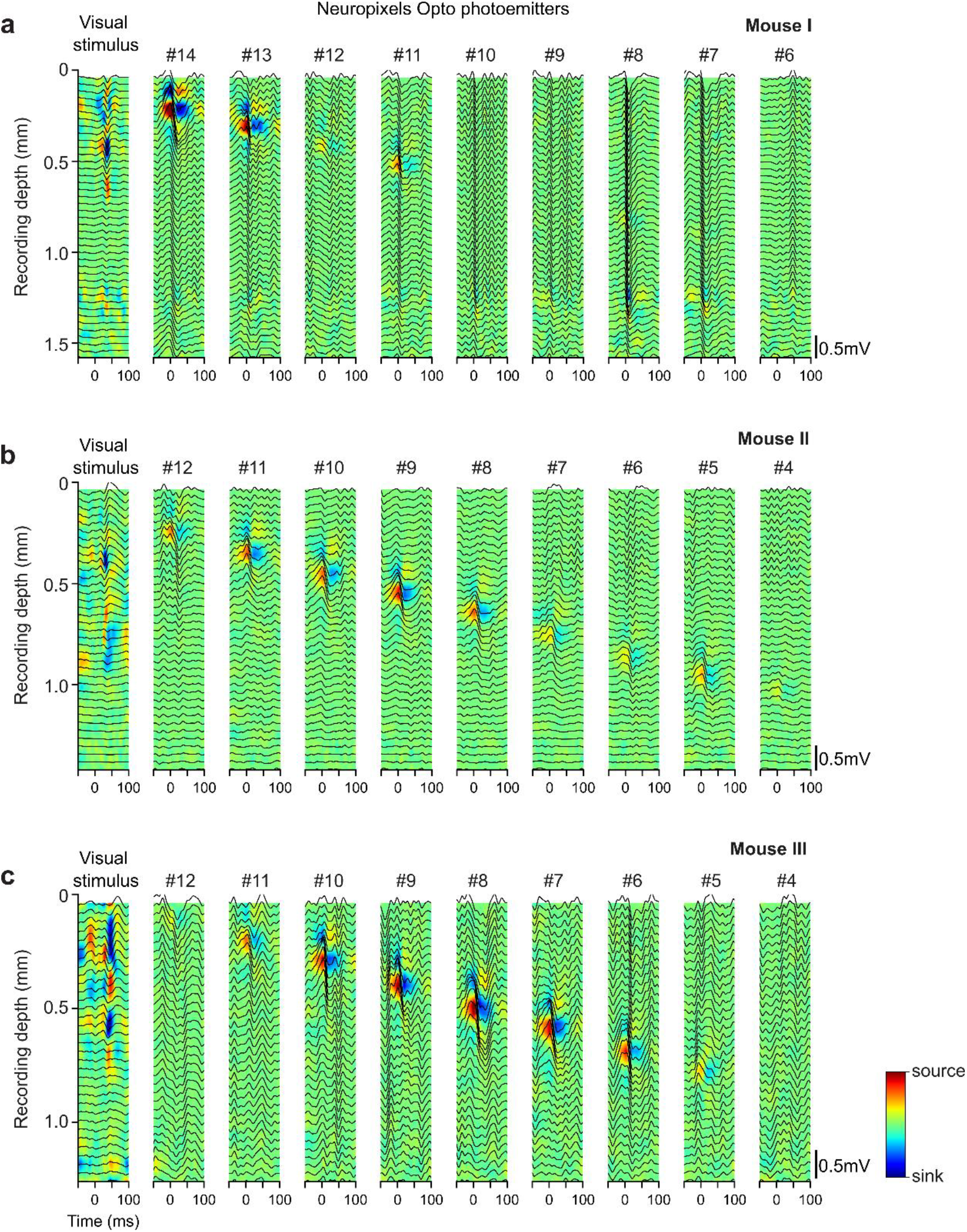
Field potentials elicited by the emitters. Local field potentials (LFPs) and current source density (CSD) profiles of activity elicited by the emitters in the primary visual cortex of mice expressing ChRmine, for the three experiments highlighted in **Fig. 3**. LFPs were computed by bandpass filtering the signal (5–60 Hz) using non-causal, zero-phase delay filtering; averaging across 40 trials; and aligning to the stimulus onsets. For visual stimulation, these onsets were contrast reversals of the checkerboard stimulus (*leftmost column*). For emitters, the onsets were relative to one of 9 emitters starting at the top of the cortex (*remaining columns*). CSD profiles show current sinks (blue, inward currents) and sources (red, outward currents), indicating neural activation specific to the emitter location. The field potentials in response to the surface laser contained artifacts and were not analyzed. **a**. Mouse I (session 1). **b**. Mouse II (session 3). **c**. Mouse III (session 3).

**Figure S10.**
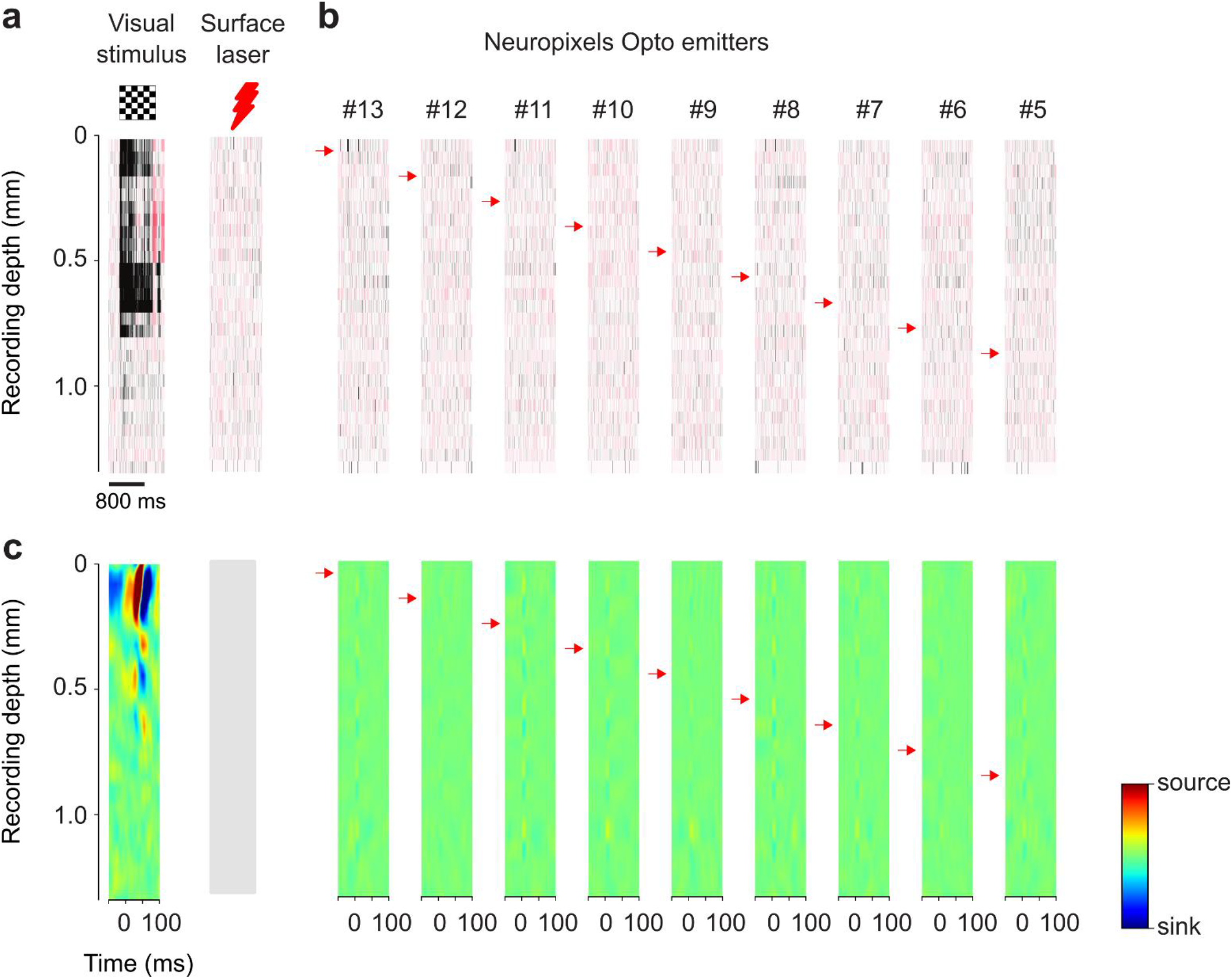
Control recordings in a mouse without opsin. The same experiment as in **Fig. 3** was conducted in a control mouse that expressed no opsin. As expected, light stimuli (external or via the emitters) did not elicit any activity. **a-b**. Firing rate as a function of depth in response to visual stimulation, external laser, and stimulating emitters Format as in **Fig. 3d-e. c**. Local field potentials and current source densities measured in the same experiments. Format as in **Fig. S9**. The field potentials in response to the surface laser (*gray rectangle*) contained artifacts and were not analyzed.

**Figure S11.**
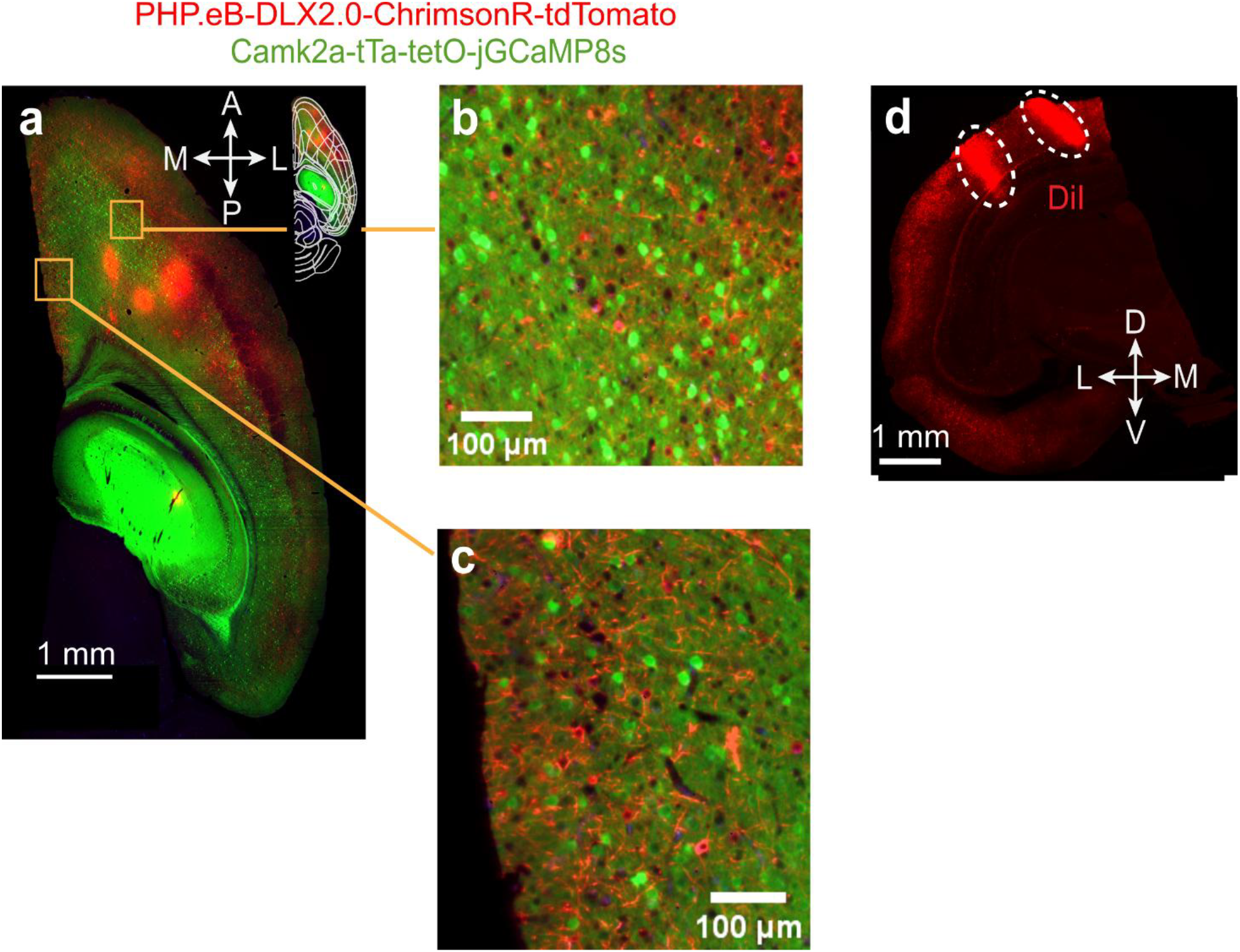
Expression of ChrimsonR-tdTomato targets neurons that do not express CaMK2a. The brain of mouse 2 (**Fig. 4**) was cleared using the SHIELD protocol (Methods). **a**. Horizontal section through the volume, showing fluorescence of ChrimsonR-tdTomato (*red*), which was targeted at inhibitory neurons via a DLX2.0 enhancer, and of GCaMP (*green*), which was targeted at excitatory neurons via a CaMK2a driver. **b-c**, Magnified regions in **a** demonstrate that tdTomato and GCAMP express in distinct neural populations. **d**. Coronal slice of the red channel showing fluorescence of ChrimsonR-tdTomato and of the fluorescent dye DiI (*dashed*), which was used to track probe trajectories. The two probe tracks traverse visual and retrosplenial cortex.

**Figure S12.**
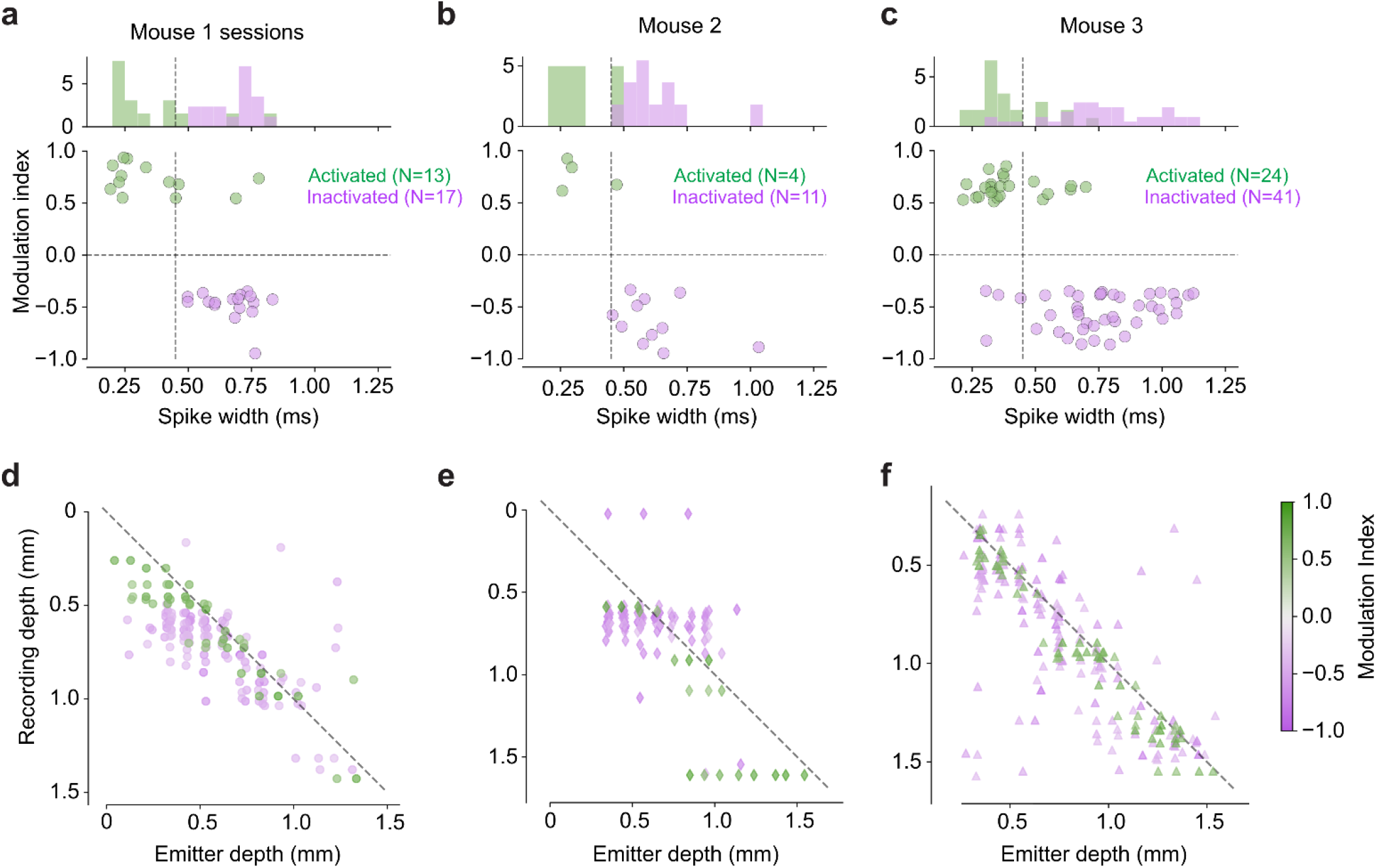
Optogenetic activation of putative inhibitory neurons in individual mice. **a-c**. Same data as in **Fig. 4e**, broken down by mouse, showing modulation index (R_1_-R_0_)/(R_1_+R_0_) (*ordinate*) as a function of trough-to-peak spike width of average waveforms. **d-f**. Same data as in **Fig. 4g**, broken down by mouse. Each panel shows the recording depth of the neurons as a function of the depth of the effective emitters. Colors indicate modulation index. Each neuron appears at one recording depth and at one or more emitter depths (if modulated by light from multiple emitters).

**Figure S13.**
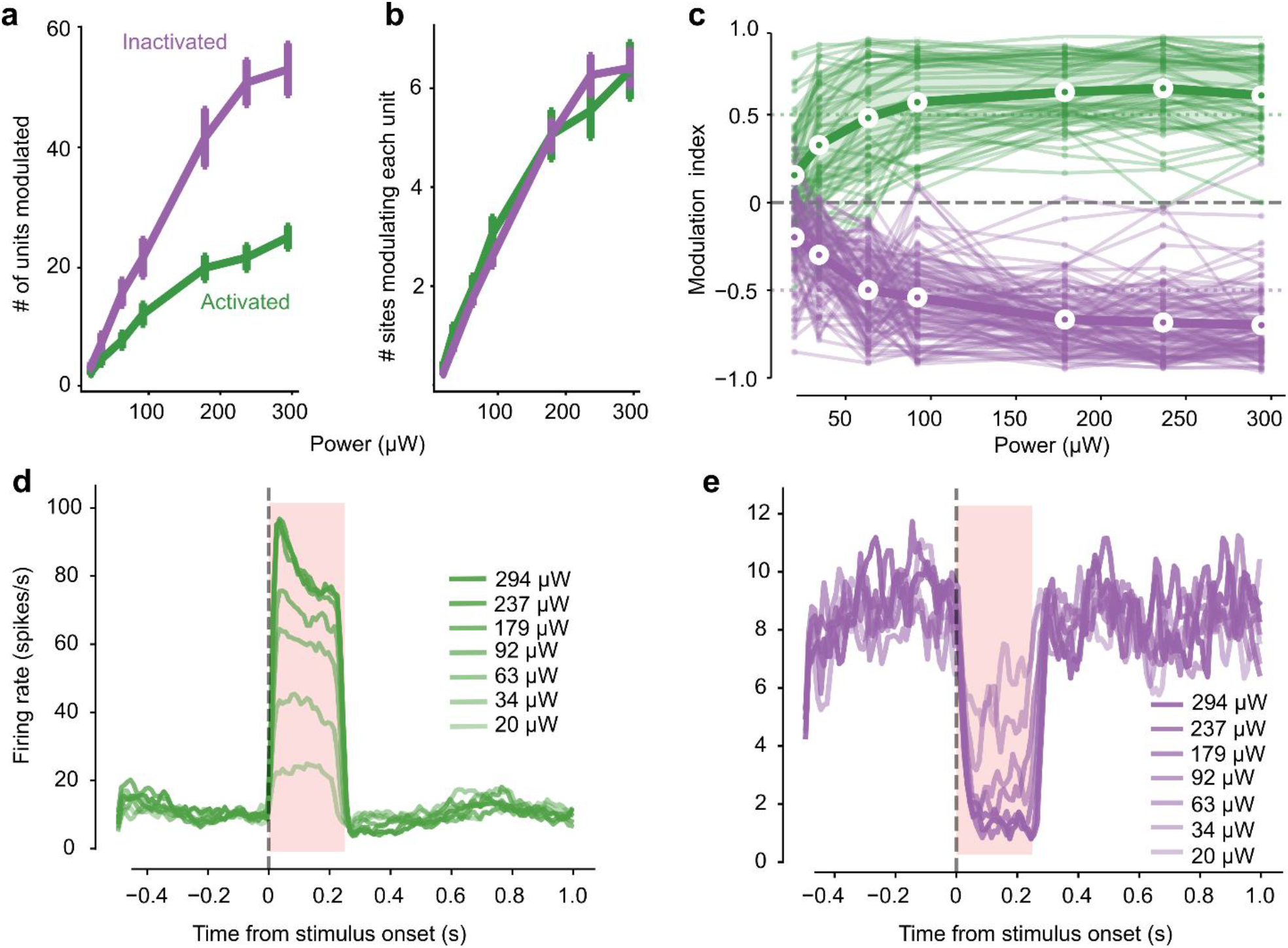
Optogenetic activation of putative inhibitory neurons: dependence on light power. Data are from an example session in mouse 3. **a**, Number of units modulated by light (same criteria as in **Fig. 4**) as a function of power of light emitted from the probe. Data in this panel and proceeding panels from a single session in mouse 3. **b**. Number of sites activating each unit at each level of light power. **c**. Modulation index as a function of laser power. All units that are eventually modulated at any power are included in the plot. **d**. Mean PSTH of activated units as a function of laser power. **e**. Same, for inactivated units.

**Figure S14.**
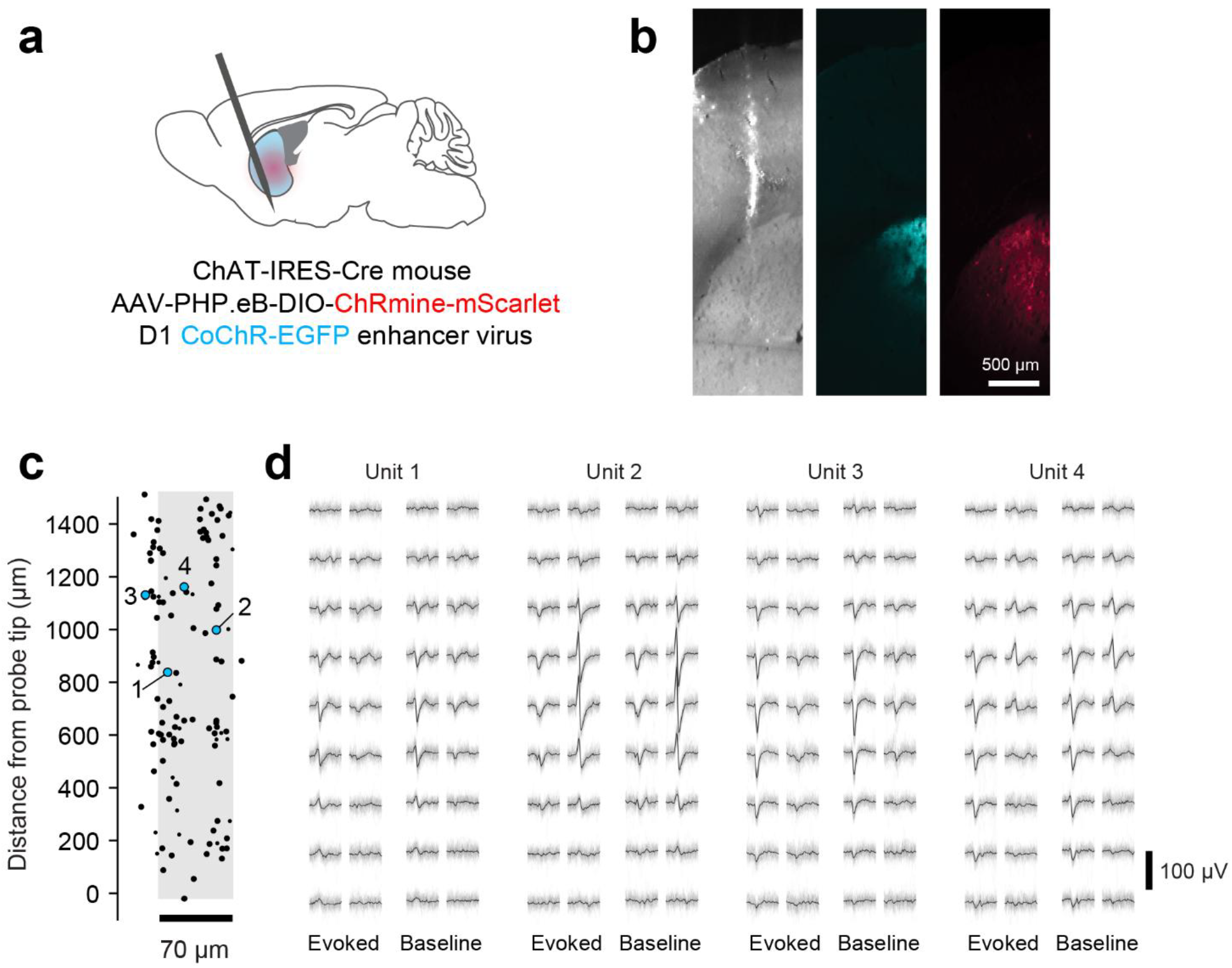
Activation of neurons by Neuropixels Opto preserves waveform shape. **a**. Schematic of reagents used in an example experiment. **b**. Virtual slice through light sheet volume of the brains used for this experiment, showing the location of the probe track and the regions of blue- or red-shifted opsin expression. **c**. Summary of unit locations within 100 µm of an emitter. Blue-tagged units appear as blue dots; there were no red-tagged units in this experiment. Large dots indicate units that pass quality metric thresholds for the complete session (ISI violations ratio < 0.5, amplitude cutoff < 0.1, presence ratio > 0.8). Small dots indicate units that pass the ISI violations ratio threshold only for the pre-stimulus baseline interval. Numbers indicate location of example units shown in panel d. **d**. Example waveforms extracted for each of four units. Each plot shows 50 randomly selected waveforms and their median (*black*), measured during the 10 ms light pulses (“Evoked”) and in the interval between trials, when no light stimulus was present (“Baseline”).

**Figure S15.**
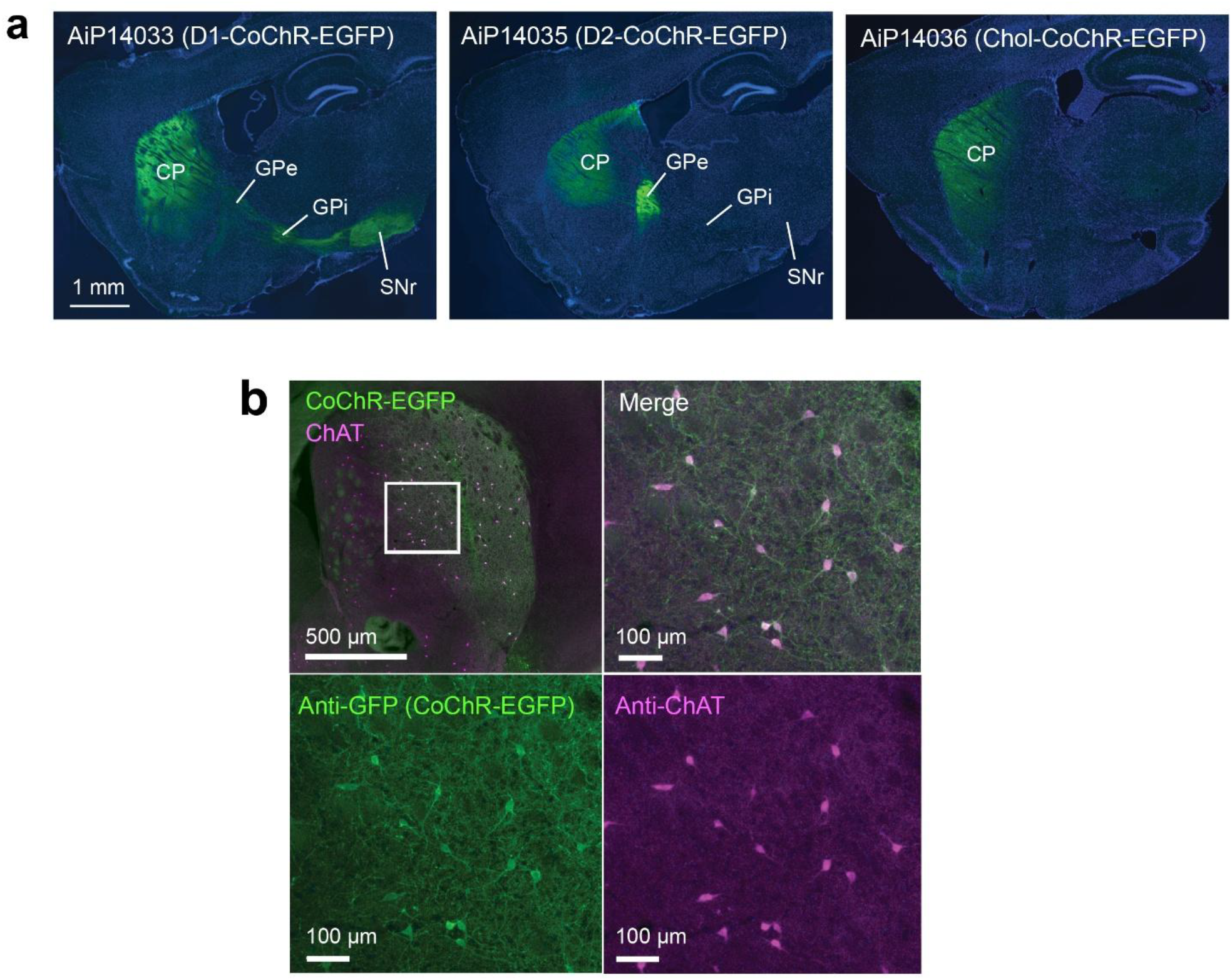
Specificity of CoChR-EGFP expression in mouse dorsal striatum. **a**, Epifluorescence image of native CoChR-EGFP expression (green) with DAPI counterstain (blue) for D1-CoChR virus AiP14033 (left), D2-CoChR virus AiP14035 (middle), and Chol-CoChR virus AiP14036 (right). Each virus was injected at 2E+9vg dose into adult C57 mice. Abbreviations: CP-caudoputamen, GPe-Globus pallidus externa, GPi-Globus pallidus interna, SNr-substantia nigra pars reticulata. **b**, High colocalization of anti-GFP (reporting CoChR-EGFP fusion protein) and anti-ChAT labeling in the dorsal striatum brain sections from AiP14036 virus-injected mouse.

**Figure S16.**
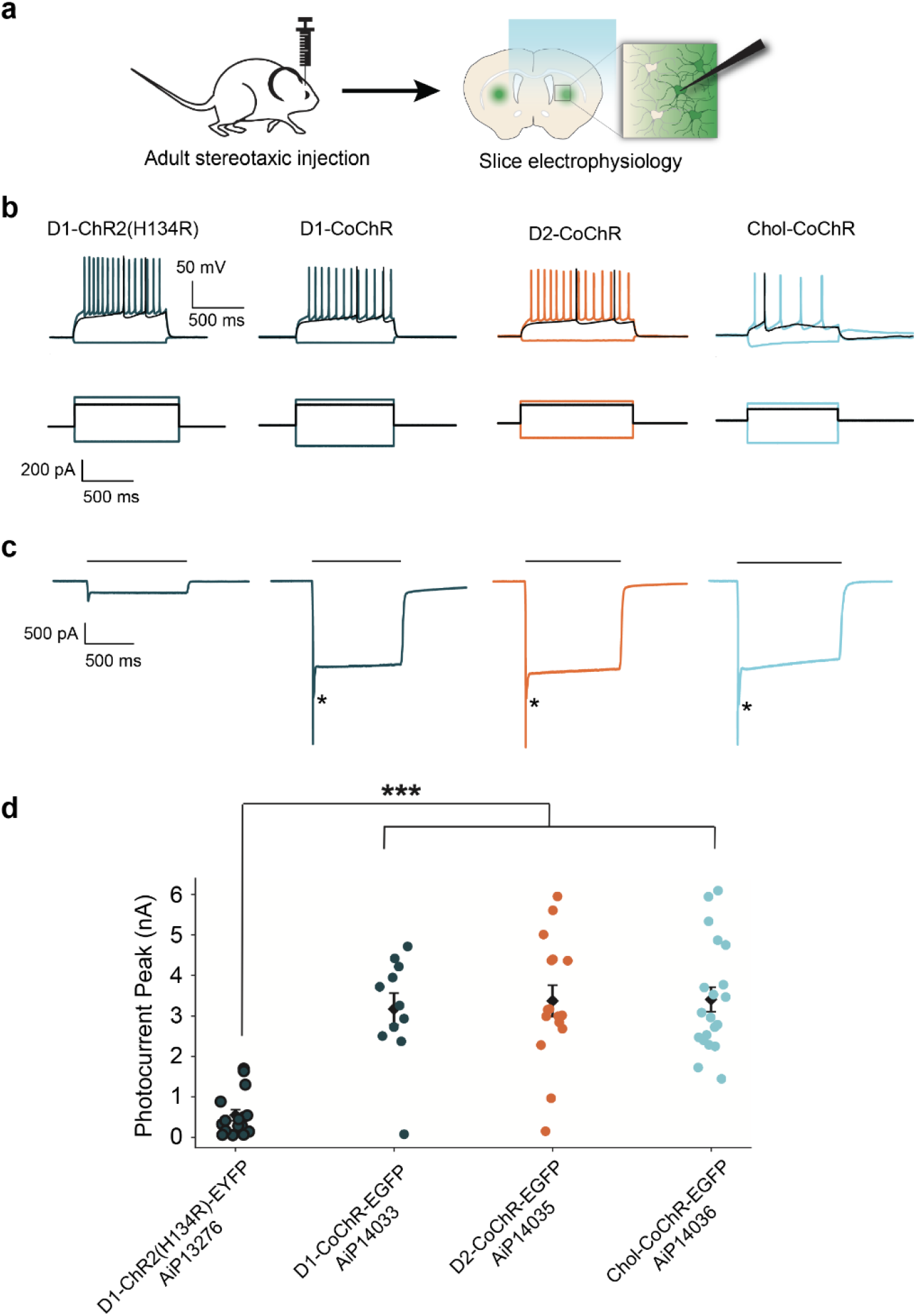
Photocurrents measured in acute brain slice patch-clamp recordings. **a**. Following the injection of AAV vectors into mouse brain (*left*) we performed acute brain slice recordings with blue light stimulation in the dorsal striatum (*right*). **b**. Whole-cell current clamp recordings in 4 representative opsin-expressing neurons, in response to a series of 1 s current injection steps (*bottom*). Putative cell type and expressed opsin are indicated. **c**. Whole-cell voltage clamp recordings from the same representative neurons, showing photocurrents evoked by 1 s blue light stimulation (*bars*). Light intensity was adjusted in each case to achieve maximal photocurrents. Asterisks denote photocurrent peak amplitude which is obscured by escape action currents observed in the CoChR-EGFP groups. **d**. Summary of peak photocurrent amplitude measurements (D1-ChR2(H134R)-EGFP, N = 20 cells; D1-CoChR-EGFP, N = 11 cells; D2-CoChR-EGFP, N = 18 cells; Chol-CoChR-EGFP, N = 21 cells). Welch’s ANOVA revealed a significant difference for mean photocurrent amplitudes across groups, and post-hoc multiple comparison test revealed that all CoChR-EGFP groups were significantly different from the D1-ChR2(H134R)-EYFP group (p < 0.0005, ***).

**Figure S17.**
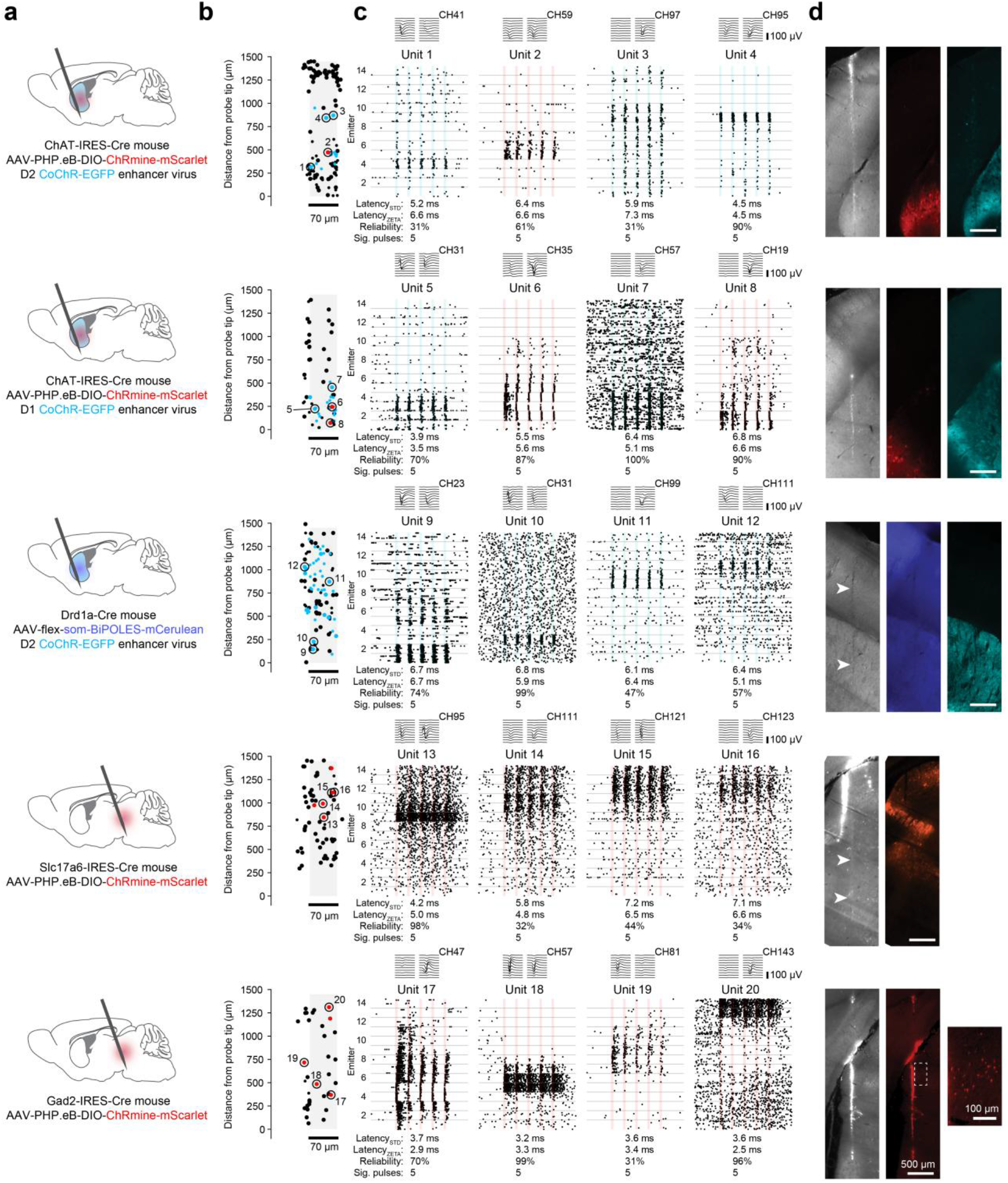
Additional examples of optotagged units. **a**. Schematic of reagents used in five different experiments. **b**. Summary of unit locations within 100 μm of an emitter. Red-tagged units appear as red dots, while blue-tagged units appear as blue dots. Untagged units appear in black. Large dots indicate units that pass quality metric thresholds for the complete session, while small dots indicate units that pass the ISI violations ratio threshold only for the pre-stimulus baseline interval. Numbers indicate location of example units shown in panel c. **c**. Stacked raster plots aligned to 20 Hz laser pulses delivered from 14 emitters, for four example units from each session. Colored vertical bars indicate the time of each laser pulse, as well as the opsin (blue-or red-shifted) that this unit expresses. If units responded to both blue and red laser presentations, they were considered to be expressing a red-shifted opsin. **d**. Virtual slices through light sheet volumes of the brains used for the same optotagging experiments shown in panels a-c, showing the location of the probe track and the regions of blue- or red-shifted opsin expression. Image colors are based on the colors shown in the schematics in panel a. White arrows indicate the probe track in cases where it is faint. In the bottom row, the expanded region shows examples of fluorescent cell bodies that are difficult to see in the zoom-out view.

**Supplementary Table 1.**
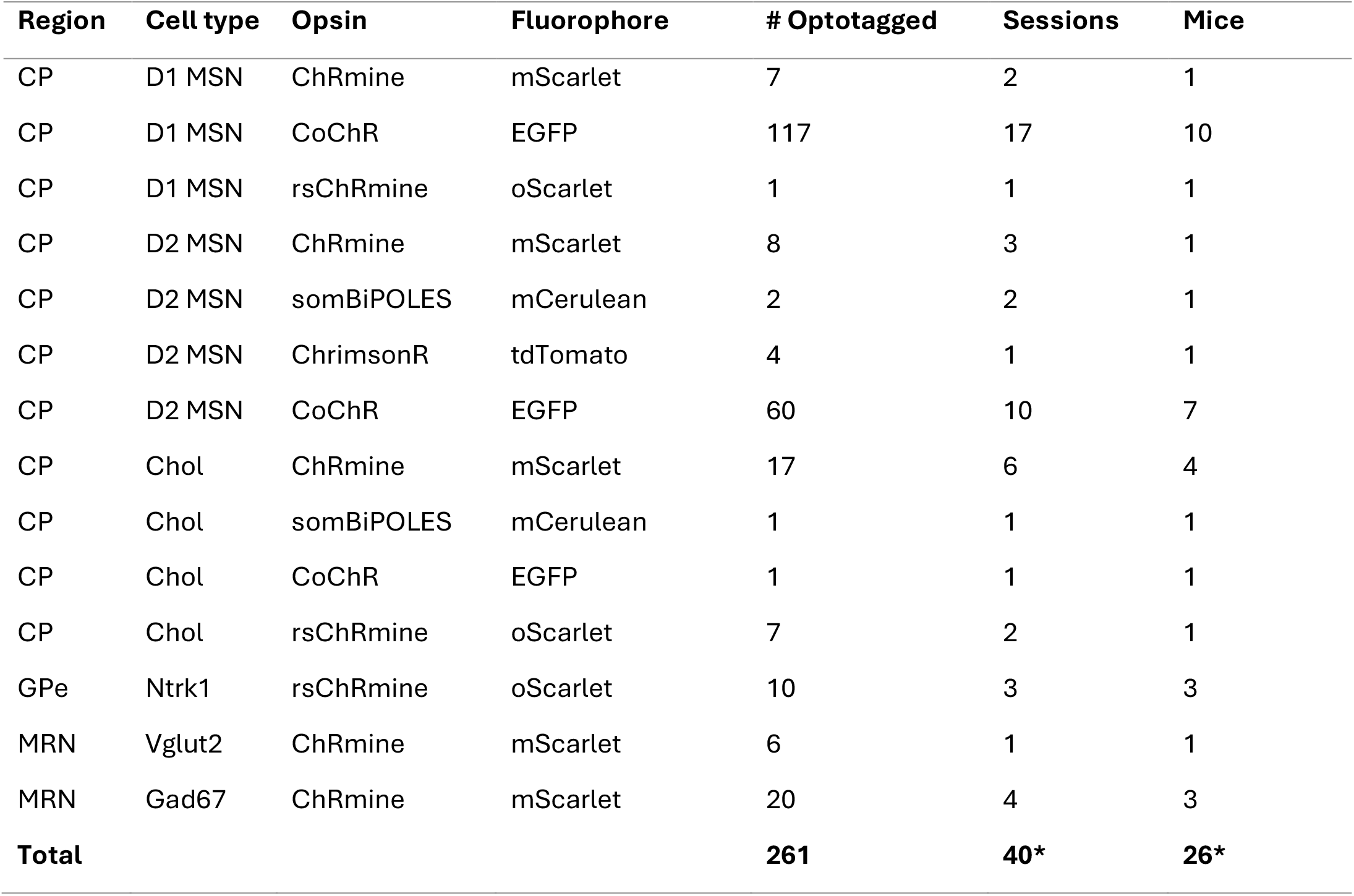
Regions and cell types for subcortical optotagging. Details of optotagged units included in **Fig. 5**. The number of units optotagged per session for a given cell type ranges between 1 and 21. Sessions may appear multiple times in this table as multiple cell types were tagged per session. * Because a session / mouse can appear in two rows, the total number of sessions and mice is in the last row is lower than the sum of the individual rows.

**Supplementary Table 2.** Device specifications for Neuropixels Opto probes. For each specification, we provide the value or range sought in 2019 before the design of the probe (“Target”) and the value obtained after designing and testing the probes in 2023 (“Result”). *The emitter switching speed indicates the physical limits (for a transition from 10% to 90%), but the software control in the current version of the probe is not real-time, so the switching speed visible to a user is much longer (multiple milliseconds).

|  | Target (2019) | Result (2023) |
| --- | --- | --- |
| Shank length | 10 mm | 10 mm |
| Shank width | 70 $\mu\text{m}$ | 70 $\mu\text{m}$ |
| Shank thickness | $\leq 35 \mu\text{m}$ | 33 $\mu\text{m}$ |
| Number of shanks | 1 | 1 |
| Silicon Base area ($W_b \times L_b$) | 7x9 mm | 9.6 x 10.2 mm |
| Package base thickness | $\leq 1 \text{ mm}$ | 1.1 mm |
| Maximum bow: base to tip | $\leq \pm 200 \mu\text{m}$ | $\leq \pm 200 \mu\text{m}$ |
| Emission wavelength (Blue) | 450-470 nm | 450 nm |
| Emission wavelength (Amber/Red) | 570-650 nm | 638 nm |
| Emitter dimension (W x L) | 15 x 35 $\mu\text{m}$ | 0.45 x 32 $\mu\text{m}$ (blue)<br>0.60 x 42 $\mu\text{m}$ (red) |
| Emitter emission patterns | Hemispherical, collimated, or focused | See Fig. 2 |
| Emitter pitch | 200 $\mu\text{m}$ | 100 $\mu\text{m}$ |
| Emitter intensity (100 $\mu\text{m}$ from shank) | 2 mW/mm<sup>2</sup> @470nm | See Fig. 2 |
| Emitter switching speed | < 1 ms | ~12 $\mu\text{s}$ * |
| Emitter count (sites/color) | 14 | 14 |
| Electrode material | TiN | TiN |
| Electrode dimension | 12 x 12 $\mu\text{m}$ | 12 x 12 $\mu\text{m}$ |
| Electrode pitch | 15 to 20 $\mu\text{m}$ | 20 $\mu\text{m}$ |
| Electrode count | 300-500 | 960 |
| Channel count | 150-300 | 384 |
| Thermal dissipation during use (difference between probe surface and tissue) | < 1°C | < 1°C |

